# Tolerance Regions for Compositional Data with Application to Reference Regions for Healthy Microbiome Profiles

**DOI:** 10.64898/2026.05.06.723285

**Authors:** Nisansala Wickramasinghe, Pankaj Choudhary

## Abstract

Imbalances in the human microbiome are associated with numerous diseases, highlighting the need for benchmarks that define healthy microbiome composition and identify abnormal deviations. Although the microbiome is increasingly studied as a potential clinical marker, statistical approaches for constructing reference regions of healthy microbiome composition remain relatively underexplored. This work develops statistical methods to construct reference regions for healthy microbiome data, addressing three main challenges. First, since microbiome data contain relative rather than absolute information, standard statistical methods are not directly appropriate. Therefore, microbiome profiles are treated as compositional data satisfying a sum constraint, and log-ratio transformations are used to analyze them in real space while preserving their relative structure. Second, reference regions are constructed as tolerance regions rather than confidence regions, so that they cover a pre-specified proportion of the healthy population with a given confidence level. The proposed framework incorporates both parametric and nonparametric approaches for constructing these tolerance regions. Parametric methods are considered when the ilr-transformed data approximately follow an elliptical distribution, where they can yield smaller regions while maintaining the desired coverage. Nonparametric approaches provide a flexible alternative by avoiding distributional assumptions. Third, because microbiome data are multidimensional and difficult to interpret, quantitative and graphical tools are introduced to assess atypicality and identify which microbial taxa contribute most to deviations from healthy profiles. Simulation studies are conducted to evaluate the performance of the proposed methods. The methodology is then demonstrated by constructing reference regions for healthy microbiome profiles using real-world data. Finally, the approach is applied to microbiome datasets comparing healthy and patient profiles to assess whether patient samples are identified as atypical and to examine which taxa contribute to these deviations. Overall, the proposed framework provides a clear and statistically robust approach for defining healthy microbiome reference regions and detecting atypical microbiome profiles.

## 1 Introduction

The human microbiome is a community of microorganisms, including bacteria, viruses, and fungi, that inhabit the human body. Considerable diversity exists in the microbiome compositions at different body sites, such as the gut, skin, mouth, and nose; and those of healthy and diseased individuals.^1^ The gut microbiome is specifically believed to play an important role in the host’s nutrient metabolism and immune system.^2^ For example, there is evidence of association between the gut microbiome composition and various health conditions, including obesity, inflammatory bowel disease, depression, and diabetes.^3,4^ Due to its potential implications for human health and well-being, microbiome research has gained significant attention recently. However, despite the growing body of literature in the field, one key question seems to have remained unanswered: What is the extent of “normal” for a healthy individual’s microbiome profile from a specific body site? The answer is expected to be a multidimensional region within which the microbiome profiles of a large proportion of the healthy population lie. Such a region accounts for the natural variation in the microbiome compositions of healthy individuals and can serve as a reference region for the composition of microorganisms inhabiting the specific body site. Clinicians can examine the consistency between an individual’s microbiome and the reference region, and a substantial deviation may indicate a health condition, making the reference region a valuable tool for diagnosis and treatment.

Indeed, reference regions, also known as “normal ranges,” for clinical markers such as blood pressure, cholesterol level, and heart rate, are widely used by clinicians as benchmarks to compare an individual’s results.^5^ For example, for blood pressure, normal is considered to be less than 120 mmHg for systolic pressure and less than 80 mmHg for diastolic pressure.^6^ Individuals with blood pressure values outside this reference region are recommended for lifestyle modifications or treatment depending on how far their values deviate from the 120/80 mmHg boundary of the region.^7^

To construct a reference region for healthy microbiome profiles, we need to address three issues. The first is that microbiome profiles are compositional in nature: they represent relative proportions rather than absolute counts. Microbiome organisms are typically grouped according to shared taxonomic ranks, with phylum, class, order, family, genus, and species being the most common ranks. In processed microbiome data, counts represent the observed abundance of each taxon in a sample. Raw counts are then normalized to reduce variability due to differences in sequencing depth across samples. This is done by dividing each taxon’s count by the total count within the sample, so that the components sum to one. This unit-sum constraint induces dependence among taxa, meaning that a change in one component necessarily affects the interpretation of the others. Hence, analyzing profiles from different samples calls for jointly considering the entire microbial community structure rather than separate taxon-wise analysis. Moreover, the traditional multivariate statistical methods, when applied directly to microbiome data, ignoring the unit sum constraint, lead to spurious results.^8,9^ A standard approach to address this issue is to treat the microbiome profiles as compositions.^10,11^ A *composition* is a vector of nonnegative real numbers that carry only relative information, meaning that information is contained only in the ratios of the components; the actual numerical values of the components are irrelevant. The components refer to parts of a whole, e.g., parts per unit (proportions), percentages, and parts per million (ppm). Although the literature on statistical tools for compositional data analysis is well developed,^9,12,^13 the problem of reference regions has not been considered. Therefore, we first propose a statistical methodology to construct a reference region for general compositional data and then apply it to the microbiome data.

The second issue is how to actually define a reference region that contains a pre-specified large fraction *p* of the population values. To fix ideas, suppose the population is univariate. In this case, by definition, the interval whose respective lower and upper limits are 100(1 − *p*)*/*2th and 100(1 + *p*)*/*2th population percentiles contain 100*p*% of the population values. However, the population distribution and hence this percentile interval are unknown and need to be estimated from the sample data. The estimated interval can be nonparametric, obtained by replacing the population quantiles with the corresponding sample order statistics, or it can be parametric. For example, under the assumption that the population distribution is Gaussian with mean *µ* and variance *σ*^2^, the percentile interval is *µ* ± *z* _{ (1+*p*)*/*2}_ *σ*, where *z*_*p*_ is the 100*p*th percentile of the standard Gaussian distribution. The estimated interval is 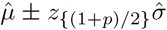, where 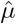 and 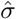 are the sample mean and standard deviation. The estimated percentile interval — whether nonparametric or parametric — is commonly taken as the reference interval in the biomedical literature.^14^ However, the fraction of population values falling in the estimated interval is not *p* anymore — it is a random quantity, because of which the estimated percentile interval is not appropriate as a reference interval.^5^ Also often used^15–17^ but not appropriate for the task is a 100*p*% *prediction interval* for a population because it is designed to capture only a single observation from the population and not a fraction *p* of the population observations.^5,18^

The appropriate reference interval is a (*p*, 1 − *α*) *tolerance interval*. It is specifically designed to contain 100*p*% of the population values with pre-specified large confidence 100(1 − *α*)%.^5,18^ In practice, it is common to take *p* ∈ {0.90, 0.95} and 1 − *α* = 0.95. By analogy with the univariate case, the appropriate reference region in the multivariate case is a (*p*, 1 − *α*) *tolerance region*. Statistical methods for constructing tolerance intervals and regions and software tools for implementing them are well developed.^19,20^ Of specific interest in this article are the nonparametric tolerance region^21^ and the classical tolerance region under the multivariate Gaussian assumption for the population distribution and its robust counterpart^22^ that is less sensitive to the potentially outlying observations that may be present in the sample data.

The third issue arises when an individual’s profile is identified as “atypical”, falling outside the reference region. Due to the multidimensional nature of the data, it is often unclear how far the profile lies from the boundary or which specific components contribute to this atypicality. To address this issue, diagnostic tools are needed to quantify the degree of atypicality and identify the components driving it.

The rest of this article is organized as follows. Section 2 reviews some basic definitions and concepts related to compositional data analysis and tolerance regions. Section 3 presents the proposed methodology for constructing a tolerance region for compositional data, detecting atypical observations, quantifying their atypicality, and generating insights into their atypicality. Section 4 describes a simulation study to evaluate the properties of the proposed methodology. Section 5 constructs the reference region for a healthy microbiome profile by applying the methodology to gut microbiome data. It then applies the approach to microbiome datasets to compare healthy and patient profiles, assess whether patient samples are identified as atypical, and examine which taxa contribute to these deviations. Section 6 concludes with a discussion. The statistical software system `R`^23^ has been used for performing the computations and data analysis in this article.

## 2 Preliminaries

### 2.1 Compositional data

Let the 1 × *D* vector **x** = [*x*_1_, …, *x*_*D*_] denote a *D*-part composition. The elements of this compositional vector are positive real numbers that represent parts of a whole and add to a constant. Without loss of generality, the constant can be assumed to be one because the numbers can be divided by their sum — an operation called *closure* — without altering the relative information contained in them. Upon closure, formally defined as

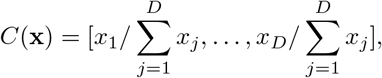

the elements of **x** can be interpreted as proportions. Thus, the sample space of compositional data, consisting of vectors of proportions adding to one, is the (*D* − 1)-dimensional unit simplex

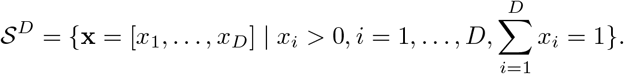

When studying a given composition, sub-compositions can be created by selecting the desired components or parts and applying the closure operation to sum them up to one. This ensures that the relative information of components within a sub-composition is consistent with the overall composition from which it is derived.

Since the compositional parts *x*_1_, …, *x*_*D*_ only contain relative information, it is common for their analysis to focus on ratios after applying a logarithmic transformation. The logratio transformation maps the data in the simplex to data in the real space, which can then be analyzed using conventional statistical methods. Several logratio transformations are available for analyzing compositional data, including the additive logratio (alr), centered logratio (clr), and isometric logratio (ilr) transformations. A detailed discussion of these transformations and their properties can be found in compositional data analysis texts,^9,12^ with additional details on alr and clr provided in the Appendix 1 of the Supplementary Document. In this article, the focus is on the ilr transformation because it maps compositions from the simplex to an unconstrained Euclidean space while preserving the Aitchison geometry, which is the natural geometry for compositional data based on logratios. This preservation is important because distances and inner products between compositions remain meaningful after transformation.

#### 2.1.1 The isometric logratio transformation

To define the ilr transformation, we follow Pawlowsky-Glahn et al.^12^ Let 𝒮^*D*^ denote the *D*-part simplex, and let **x** = [*x*_1_, …, *x*_*D*_] ∈ 𝒮^*D*^ be a composition. For **x, y** ∈ 𝒮^*D*^, the Aitchison inner product is defined as

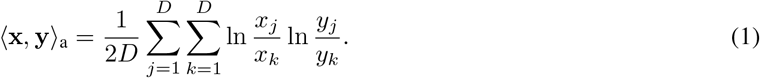

Let {**e**_1_, …, **e**_*D*−1_} be an orthonormal basis for 𝒮^*D*^ with respect to the Aitchison inner product. Then any composition **x** ∈ 𝒮^*D*^ can be represented as

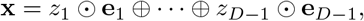

where ⊕ and ⊙ denote perturbation and powering, respectively. The coefficients

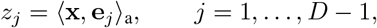

provide the coordinates of **x** with respect to the orthonormal basis. Let **z** = [*z*_1_, …, *z*_*D*−1_] be the vector of these coordinates. The ilr transformation of **x** is defined as ilr(**x**) = **z**. Thus, ilr maps 𝒮^*D*^ into ℝ^*D*−1^, giving the coordinates of **x** with respect to the chosen orthonormal basis. Next, let **Ψ** be a (*D* − 1) × *D* matrix whose *j*th row contains the clr coordinates of **e**_*j*_, *j* = 1, …, *D* − 1. Then, each row of **Ψ** adds to zero. In addition, it can be seen that **z** = ln(**x**)**Ψ**^′^, implying that the ilr coordinates are *logcontrasts* — linear combinations of ln *x*_1_, …, ln *x*_*D*_ with coefficients adding to zero. For this reason, **Ψ** is also called the *contrast matrix* associated with the basis. The inverse ilr transformation is **x** = ilr^−1^(**z**) = *C*(exp(**zΨ**)).

Since an orthonormal basis is not unique, it can be constructed in specific ways to enhance the interpretability of the associated ilr coordinates. One well-known construction leads to the following coordinates called *balances*:^12^

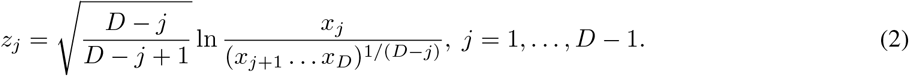

Here *z*_*j*_ is a normalized logratio capturing the relative information between part *j* and the geometric mean of the group of parts that come after *j*. Depending on the sign of *z*_*j*_, the part *j* has more or less weight in the composition than the denominator group.

The alr, clr, and ilr transformations are linearly related,^24^ but they differ in properties that are important for statistical modeling. The alr transformation maps compositions into *D* − 1 coordinates, but these coordinates depend on the choice of denominator component and do not preserve Aitchison distances in the simplex. The clr transformation preserves Aitchison geometry, but it represents a *D*-part composition using *D* linearly dependent coordinates. Consequently, the covariance matrix of clr-transformed data is singular, and probability models specified directly for clr coordinates are degenerate in ℝ^*D*^. In contrast, the ilr transformation represents a *D*-part composition using *D* − 1 unconstrained coordinates defined with respect to an orthonormal basis. Thus, the ilr transformation preserves Aitchison geometry while avoiding other limitations of the alr and clr transformations. For this reason, the methodology in this article is developed using ilr coordinates.

#### 2.1.2 Compositional approach to statistical modeling

Let **X** be a random composition in 𝒮^*D*^. Consider an orthonormal basis of 𝒮^*D*^ whose associated contrast matrix is **Ψ**. Let **Z** = ilr(**X**) = ln(**X**)**Ψ**^′^ be the random vector in ℝ^*D*−1^, representing the ilr coordinate vector of **X** with respect the given basis. Let *B* be an event in ℝ^*D*−1^ and *A* = ilr^−1^(*B*) be the corresponding event in 𝒮^*D*^. Then, from the *principle of working on coordinates*,^12^ it follows that

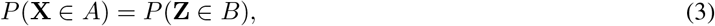

and the probability in the simplex does not depend on the chosen basis and the corresponding **Ψ**. As a result, properties of random compositions can be studied using their transformed ilr coordinates for which traditional statistical techniques are available and back transforming the results to the simplex. Here we assume that **Z** and hence **X** follow continuous multivariate probability distributions.

### 2.2 Tolerance regions

Consider a *d*-variate population (*d* ≥ 1) and let **Y**_1_, **Y**_2_, …, **Y**_*n*_ denote a random sample from this population. Let **Y** denote a future observation from this population in that it is independent of the sample data. Next, let the region *T* = *T* (**Y**_1_, **Y**_2_, …, **Y**_*n*_) be a function of sample data representing a subset of the sample space of the population. The probability content of region *T* conditional on the sample data, given as *P*_**Y**_(**Y** ∈ *T* | **Y**_1_, …, **Y**_*n*_), is the fraction of observations from the population that are contained within the region. Because this fraction is a function of the sample data, it is a random quantity and has a sampling distribution. For a specified proportion *p* and level of confidence 1 − *α*, the region *T* is called a 100(*p*, 1 − *α*)% *tolerance region* for the population of **Y** if

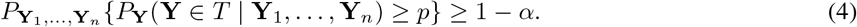

Thus, the tolerance region *T* is designed to contain at least 100*p*% of the observations from the sampled population with 100(1 − *α*)% confidence.

#### 2.2.1 Univariate normal tolerance intervals

Taking *d* = 1 and considering *T* as an interval whose limits are functions of the sample data gives a tolerance interval. Suppose the population follows a 𝒩_1_(*µ, σ*^2^) distribution with mean *µ* and variance *σ*^2^. In this case, for *n >* 1, the classical two-sided tolerance limits^19^ are of the form 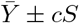, where 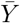 is the sample mean, *S* is the sample standard deviation, and the tolerance factor *c* = *c*(*p, α, n*) is determined such that the probability statement (4) holds. Although this *c* is not available in a closed form, several approximations are available.^19^ Next, we consider a generalization to the multivariate case.

#### 2.2.2 Multivariate normal tolerance regions

Suppose the population follows a 𝒩_*d*_(***µ*, Σ**) distribution with mean vector ***µ*** and nonsingular covariance matrix **Σ**. Assuming *n > d*, let 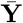 and **S**_*y*_ respectively denote the sample mean vector and sample covariance matrix of the data. Also let

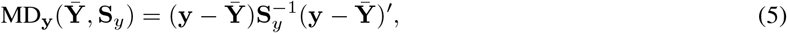

denote the squared Mahalanobis distance between a point **y** and the center 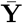 with respect to the dispersion matrix **S**_*y*_. The classical tolerance region is an ellipsoid of the form^19^

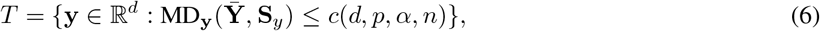

where the tolerance factor *c*(*d, p, α, n*) is determined to satisfy the probability requirement

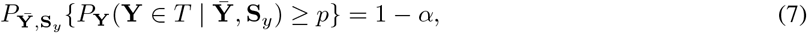

which is equivalent to (4) when coverage probability is taken to be 1 − *α*. The tolerance factor represents the squared distance between the center and the ellipsoid boundary. As in the univariate case, *c* is not available in a closed form, but several approximations are available.^19,25,^26 Of specific interest in this article is a Monte Carlo approximation. To describe it, let

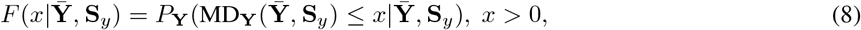

denote the cumulative distribution function (cdf) of the squared Mahalanobis distance (5) conditional on 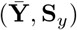. We can write (7) as

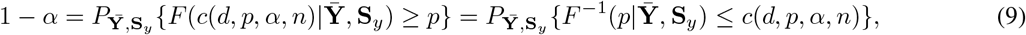

implying that *c* is the (1 − *α*)th quantile of the sampling distribution of 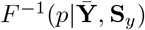. It is well known that the sampling distribution of 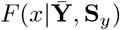 and hence that of its inverse 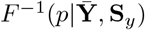 and also the tolerance factor *c* do not depend on (***µ*, Σ**).^19^ This assertion follows from a Proposition 1 given in Section 2.3. Thus, *c* can be computed using a Monte Carlo approach by setting (***µ*, Σ**) = (**0, I**), where **I** is the identity matrix. The algorithm for more general case is given in Section 2.3.

#### 2.2.3 Multivariate normal robust tolerance regions

The estimator 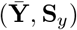 of (***µ*, Σ**) used to construct the classical tolerance region is not robust to outlying observations in the data. In such a scenario, the lower bound *p* on the probability content in (7) cannot be relied upon.^22^ To address this issue, robust tolerance regions^22^ have been developed. These are obtained by computing a robust estimator (**M**_*y*_, **V**_*y*_) of (***µ*, Σ**) from the sample data and replacing (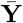, **S**_*y*_) in (6) with (**M**_*y*_, **V**_*y*_) so that

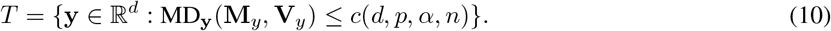

As before, the tolerance factor *c*(*d, p, α, n*) is chosen to satisfy the requirement

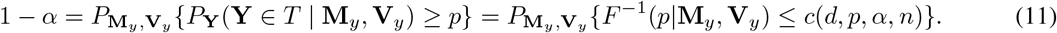

Although a variety of robust estimators of ***µ*** and **Σ** are available in the literature,^27^ the *affine equivariant* estimators, i.e., those that satisfy

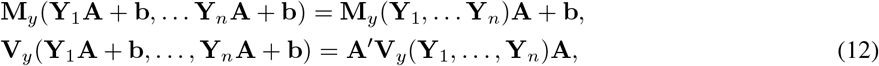

for any vector **b** and nonsingular matrix **A**, are especially appealing for constructing tolerance regions. This is because with affine equivariant estimators, the probability on the right in (11) does not depend on (***µ*, Σ**).^22^ As a result, the tolerance factor *c* can be readily computed by modifying the Monte Carlo approach for computing *c*;^22^ see Algorithm 1.

Following Boente and Farall,^22^ we consider three robust estimators of ***µ*** and **Σ**. The first is the minimum covariance determinant (MCD) estimator introduced by Rousseeuw.^28^ It is obtained by identifying a subset of the data points whose covariance matrix has the smallest determinant and taking the mean and a scaled covariance matrix of the subset. The second estimator is the Stahel-Donoho estimator (SD) introduced by Stahel^29^ and Donoho^30^ and popularized by Maronna and Yohai.^31^ It involves taking a weighted mean and scaled weighted covariance matrix of the data, where each sample observation is weighted according to a measure of its outlyingness. The third estimator is the S-estimator studied by Davies.^32^ It is obtained by minimizing a scalar measure of dispersion of the data cloud subject to a constraint that restricts the influence of the outlying data points. These estimators are described in detail elsewhere^27^ and have been implemented in the `R` package `rrcov`.^33^

### 2.3 Tolerance regions for elliptical distributions

In practice, oftentimes a more flexible distribution than normal is needed to adequately model the data. One important class of such distributions is *elliptical distributions*.^34^ In addition to the normal distribution, its members include the *t* and contaminated normal distributions that allow modeling of data with heavier tails than normal. A random vector **Y** follows an elliptical distribution with location vector ***µ*** and nonsingular dispersion matrix **Λ** if its density is

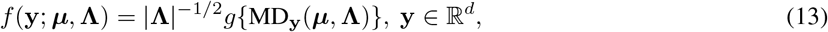

where MD_**y**_(***µ*, Λ**) = (**y** − ***µ***)**Λ**^−1^(**y** − ***µ***)^′^ is the squared Mahalanobis distance given by (5) and *g*(·) ≥ 0 is a density function satisfying ∫*g*(**yy**^′^)*d***y** = 1. Since the density (13) is constant on concentric ellipsoids MD_**y**_(***µ*, Λ**) = *k, k >* 0, the distribution is also known as an *elliptically contoured distribution*. If **C** is a nonsingular matrix such that **CΛ**^−1^**C**^′^ = **I** and **U** is a random vector with density *g*, then **Y** has the stochastic representation **Y** = **UC** + ***µ***. Such a **C** can be obtained, e.g., using the Cholesky or spectral decompositions of **Λ**. If the mean exists, ***µ*** is the mean of **Y** and if covariances exist, the covariance matrix **Σ** is proportional to **Λ**.

Despite the widespread use of the elliptical distributions for data analysis, the problem of constructing tolerance regions has not been considered for a nonnormal member of the class. Here we proceed along the lines of the methodology for the normal case to develop a general methodology that is applicable for any elliptical distribution.

Based on a random sample **Y**_1_, …, **Y**_*n*_ from the population (13) with *n > d*, let 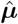 be the estimator of ***µ*** and the nonsingular dispersion matrix 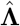 be the estimator of **Λ**, obtained, e.g., using the method of maximum likelihood (ML). If the model (13) has additional parameters besides (***µ*, Λ**), e.g., the degrees of freedom in case of a *t* distribution, these are assumed to be known. Like the normal case, the tolerance region for the general case is an ellipsoid of the form (6):

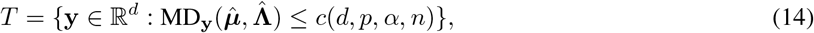

where the tolerance factor *c* is chosen to satisfy the requirement

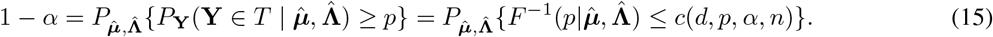

Here *F* is the conditional cdf of MD_**Y**_ given 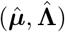. Thus, as in 9, *c* is the (1−*α*)th quantile of the sampling distribution of 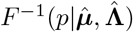.

#### Proposition 1.

*Suppose the estimators* 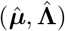 *of* (***µ*, Λ**) *are affine equivariant. Then, the sampling distribution of* 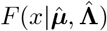, *x >* 0 *and hence that of* 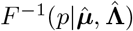, 0 *< p <* 1 *do not depend on* (***µ*, Λ**).

The proof is given in Appendix 2 of the Supplementary Document. This result also holds for the univariate case (*d* = 1). It follows from the result that when affine equivariant estimators are used, *c* can be computed using a Monte Carlo approach by setting (***µ*, Λ**) = (**0, I**); see Algorithm 1. Moreover, the assertions made in Section 2.2 regarding *F, F* ^−1^ and tolerance factors for normal distribution being free of unknown parameters follow from this result by taking **Λ** = **Σ**. This is because the estimators (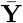, **S**_*y*_) in the classical case and (**M**_*y*_, **V**_*y*_) in the robust case are affine equivariant. The next result establishes the equivariance property for ML estimators of (***µ*, Λ**) under any elliptical distribution, allowing the use of Proposition 1 to compute *c* with ML estimators.

#### Proposition 2.

*The ML estimators* 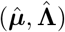 *of* (***µ*, Λ**) *in the elliptical distribution model* (13) *are affine equivariant*.

The proof is given in Appendix 2 of the Supplementary Document. The elliptical distribution of specific interest in this article is the multivariate *t* distribution, denoted as **Y** ~ *t*_*d*_(***µ*, Λ**, *ν*), where *ν* is the degrees of freedom. Its density is

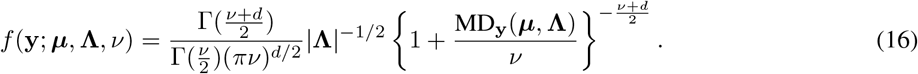

Its mean exists for *ν >* 1 and covariances exist for *ν >* 2, with **Σ** = {*ν/*(*ν* − 2)} **Λ**. The parameters of this model can be estimated by the ML method using the expectation-maximization (EM) algorithm and its extensions.^35^ We use the `R` package `fitHeavyTail`^36^ for this task. Although *ν* can be also be treated as an unknown parameter, but for modeling in practice, it is often the values such as 5 or 10 that are of interest. So we treat *ν* as a known parameter and use goodness-of-fit considerations to determine its value.

A Monte Carlo algorithm for computing tolerance factor *c* is as follows:

#### Algorithm 1.

**Tolerance factor**

For given (*d, p, α, n*) and large values for numbers of replication *N*_1_ and *N*_2_:

1. Generate *N*_1_ independent draws 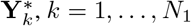 from the distribution (13) with (***µ*, Λ**) = (**0, I**)
2. Generate a random sample **Y**_*i*_, *i* = 1, …, *n* from the distribution (13) with (***µ*, Λ**) = (**0, I**) and compute the estimates 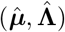
3. Calculate the squared distances 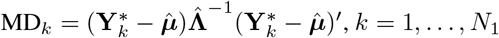
4. Compute the *p*th sample quantile of 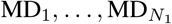; call it 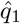
5. Repeat Steps 2-4 *N*_2_ times to get 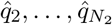
6. Compute the (1 − *α*)th sample quantile of 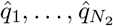 and take it as *c*(*d, p, α, n*)

When **Y** ~ 𝒩_*d*_(***µ*, Σ**), **Σ** ≡ **Λ**, and depending upon whether the classical or robust region is required, (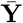, **S**_*y*_) or (**M**_*y*_, **V**_*y*_) is used as 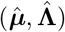 in Algorithm 1. Likewise, when **Y** ~ *t*_*d*_(***µ*, Λ**, *ν*), *ν* is known and the ML estimators 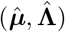 are used. In this work, we have set (*N*_1_, *N*_2_) = (10^4^, 10^4^).

### 2.4 Nonparametric tolerance regions

Nonparametric tolerance regions provide a robust method for constructing tolerance regions directly from sample data — given that the sample size requirement is met. This approach is particularly useful when dealing with continuous data that does not fit into a parametric model or when constructing a tolerance region for a parametric model is challenging. However, it is important to note that for a given sample size, it is possible that a nonparametric tolerance region satisfying specific content and coverage requirements may not exist.

The 100(*p*, 1 − *α*)% nonparametric tolerance region *T* is constructed to satisfy (4). When *T* is the union of *k* (1 *< k < n*) equivalent blocks, the conditional probability *P*_**Y**_(**Y** ∈ *T* | **Y**_1_, …, **Y**_*n*_) follows a Beta distribution with parameters *k* and *n* − *k* + 1.^37^ In this case, *k* is chosen to satisfy (4), which simplifies to the following:

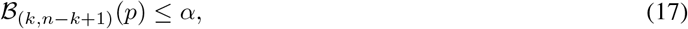

We follow the method outlined in Liu et al.^21^ for constructing distribution-free hyperrectangular tolerance regions. Initially, *n* + 1 equivalent blocks are generated from the sample **Y** as *B*_1_, …, *B*_*n*+1_. The next step involves determining the minimum number of blocks, denoted as *k*_0_, that meet (17) for given values of *n, p*, and 1 − *α*. The 100(*p*, 1 − *α*)% nonparametric tolerance region is then given by the union of the last *k*_0_ blocks: 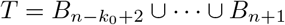.

However, due to the discrete nature of *k*_0_ and *n*, the actual confidence level of the tolerance region can exceed the intended 1 − *α*. This implies that the tolerance region may cover a larger proportion of the population distribution than the specified target coverage, which is consistent with the definition (4). Additionally, *T* forms a two-sided *d*-dimensional tolerance hyperrectangle only if *k*_0_ ≤ *n* − (2*d* − 1). The existence of such a *k*_0_ is guaranteed only if the *n* is large enough to satisfy

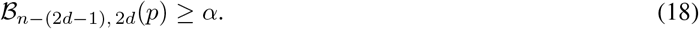

## 3 Tolerance regions for compositional data

Let **X** denote a *D*-part random composition in 𝒮 ^*D*^ representing the population of interest. Suppose the data consists of a random sample of *n* (≥ *D*) observations **X**_1_, …, **X**_*n*_ from this population. These are 1 × *D* vectors. We would like to use the sample data to construct a 100(*p*, 1 − *α*)% tolerance region for **X**. Let **Z** = ilr(**X**) and **Z**_*i*_ = ilr(**X**_*i*_) respectively denote the 1 × (*D* − 1) vectors of ilr coordinates of **X** and **X**_*i*_, *i* = 1, …, *n* with respect to a given orthonormal basis for 𝒮 ^*D*^ that has **Ψ** as the associated (*D* − 1) × *D* contrast matrix. The ilr coordinates **Z**_1_, …, **Z**_*n*_ are a random sample from the population represented by **Z**.

### 3.1 Tolerance regions for X

To get a tolerance region for **X** in 𝒮^*D*^, we first construct a tolerance region for its ilr coordinate **Z** in ℝ^*D*−1^. Identifying (*d*, **Y**) in Section 2.2 with (*D* − 1, **Z**), we get *T* as a 100(*p*, 1 − *α*)% tolerance region for **Z**. The region can be parametric or nonparametric. Inverting *T* gives a tolerance region *R* for **X**, as shown by the following result.

#### Proposition 3.

*For a given orthonormal basis for* 𝒮^*D*^ *and* **Ψ** *as the associated contrast matrix, consider a* 100(*p*, 1 − *α*)% *tolerance region T for the ilr coordinate* **Z** *of* **X**. *Upon inverting it by applying the inverse ilr transformation, we get the corresponding* 100(*p*, 1 − *α*)% *tolerance region for* **X** *in* 𝒮^*D*^ *as:*

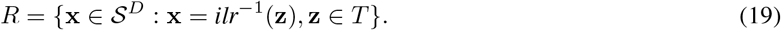

*This simplicial tolerance region is invariant to the chosen basis and the associated* **Ψ**.

Its proof is omitted as it follows directly from the principle of working on coordinates^12^ (see Section 2.1.2). When both *p* and 1 − *α* are large, e.g., (*p*, 1 − *α*) = (0.95, 0.95), the regions *R* and *T* can respectively serve as reference regions for compositional data and their ilr coordinates. Because of the correspondence between the tolerance regions for **X** and **Z** given by Proposition 3, to determine whether or not an individual’s composition falls outside the reference region of **X**, it suffices to determine whether or not its ilr coordinate falls outside the reference region of **Z**. When a composition or equivalently its coordinate fall outside the reference region, the composition may be considered atypical, potentially flagging the individual for additional investigations.

### 3.2 A *p*-index to quantify atypicality of a composition

When a composition is deemed atypical, quantifying the degree of its atypicality is essential for follow-up investigations. For this task, although one can measure how far the observation falls from the tolerance region boundary, the unbounded nature of this distance makes it hard to interpret it as a measure of atypicality. Here, we develop a probability-based index that is easier to interpret.

To describe the new index, let **x**_0_ be the composition of a given individual and **z**_0_ be its ilr coordinate. Consider a region of the same form as the initial 100(*p*, 1 − *α*)% tolerance region that passes through the point **z**_0_ and find the probability content *p*_0_ so that the region can be interpreted as a 100(*p*_0_, 1 − *α*)% tolerance region in ℝ^*D*−1^ for **Z**. The point **x**_0_ would fall on the boundary of the corresponding 100(*p*_0_, 1 − *α*)% tolerance region for **X** in 𝒮^*D*^ obtained via Proposition 3. We refer to the probability content *p*_0_ as the *p-index* of the observation **x**_0_. The *p*-index can also be interpreted as the smallest probability content for which the observation would not fall outside the corresponding tolerance region and hence will not be considered atypical. Comparing the *p*-index of a point with the probability content *p* of the initial 100(*p*, 1 − *α*)% tolerance region, we see that the *p*-index is at most *p* for the points on the boundary of the tolerance region and those inside it, whereas it is greater than *p* for the points outside. Thus, for an observation outside the reference region, its *p*-index can effectively quantify the extent of the observation’s atypicality.

In the parametric case wherein an ellipsoidal tolerance region based on an elliptical distribution is used, the *p*-index of an observation is related to its squared Mahalanobis distance from the center. Let *c*_0_ be the distance of **z**_0_ from the data center. Since **z**_0_ is a point on the boundary of the new tolerance region, the corresponding tolerance factor equals *c*_0_. As the observation moves further away from the center, whether in ℝ^*D*−1^ or 𝒮^*D*^, its *p*-index increases from zero to one. The index equals zero for an observation that falls right at the center — the “most typical” point. On the other extreme, the index equals one in the limit as the observation moves further away from the bulk of the data. Thus, as the *p*-index of an observation increases, its atypicality relative to the observed data increases. Next, we present a result that provides a way to compute it.

#### Proposition 4.

*Consider the set up of Proposition 1. The p-index of an observation equals the αth quantile of the sampling distribution of* 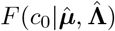.

The proof is given in Appendix 2 of the Supplementary Document. Thus, in case of an elliptical distribution, the *p*-index can be easily computed using a Monte Carlo approach by taking (***µ*, Σ**) = (**0, I**) without any loss of generality.

#### Algorithm 2.

***p*-index in case of an elliptical distribution**

For given (*d, α, n, c*_0_) and large values for *N*_1_ and *N*_2_:

1. Generate *N*_1_ independent draws 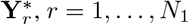 from the distribution (13) with (***µ*, Λ**) = (**0, I**)
2. Generate a random sample **Y**_*i*_, *i* = 1, …, *n* from the distribution (13) with (***µ*, Λ**) = (**0, I**) and compute the estimates 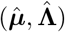
3. Calculate the squared distances 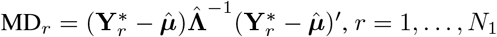
4. Compute the proportion of 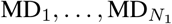 that are less than or equal to *c*_0_; call it 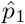
5. Repeat Steps 2-4 *N*_2_ times to get 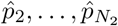
6. Compute the *α*th sample quantile of 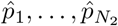 and take it as the *p*-index

In nonparametric case, the *p*-index can be calculated directly by determining the minimum number of equivalent blocks needed to form a two sided hyperrectangular tolerance region, assuming that **z**_0_ is located on the boundary. Let us denote this number as 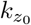. Then, the *p*-index of an observation equals the *α*th quantile of the beta distribution with parameters 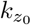 and 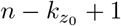.

### 3.3 Generating insights into atypicality of a composition

For a composition deemed atypical, it is helpful to determine the components of the composition that may have contributed to its atypicality. This task requires making inferences about individual components and visualizing them, which present challenges because the compositional data are multivariate and contain only relative information. To address this issue, we adapt the visualization tools^38^ originally developed for interpreting multivariate outliers in compositional data. Their usage is illustrated in Section 5.

The tools rely on *D* variables 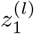, *l* = 1, …, *D* — called *ilr variables*,^38^ which are constructed as follows. Recall that an orthonormal basis for 𝒮^*D*^ can be chosen to represent a compositional vector **x** = [*x*_1_, …, *x*_*D*_] using the ilr coordinate vector **z** = [*z*_1_, …, *z*_*D*−1_] given by (2). The first coordinate *z*_1_ captures all the information contained in *x*_1_ relative to the remaining *D* − 1 components. Take 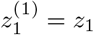. Next, rearrange the components so that *x*_2_ becomes the first component in the composition, apply the ilr transformation (2), and take the first coordinate as 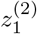. Repeat this process so that in the end we have *D* variables 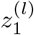, *l* = 1, …, *D*, wherein 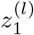 captures all the information in *x*_*l*_ relative to the remaining *D* − 1 components. More precisely, for *l >* 2, 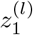 is obtained by permuting the components of **x** as (*x*_*l*_, *x*_1_, …, *x*_*l*−1_, *x*_*l*+1_, …, *x*_*D*_), applying (2), and taking the first coordinate. Thus,

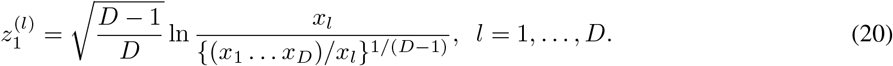

Together, these *D* variables capture all the univariate relative information about each individual component compared to the remaining components.

The data on ilr variables are plotted as a *parallel plot*^38^ to get a visual representation of the multivariate structure of the observations. In a parallel plot, multiple axes representing ilr variables are drawn parallel to each other. The observations are depicted as lines that connect the values of the ilr variables from the same individuals. The patterns and intersections of the lines provide insights into the multivariate structure of the data. Furthermore, the unusual data points often stand out as lines that deviate substantially from the others.

For an atypical observation, the *uniplots*^38^ of ilr variables can be used to visualize the parts of the composition that may be contributing to its atypicality. Uniplots are a collection of *D* scatterplots — one scatterplot for each ilr variable in which the vertical axis represents the value of ilr variable and the horizontal axis represents the observation number. We propose to superimpose on each scatterplot two horizontal lines representing the univariate tolerance limits for the corresponding ilr variable. The 100(*p*, 1 − *α*)% univariate tolerance interval for each ilr random variable 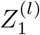 is an interval (*L, U*), where the lower and upper limits are computed using the univariate counterparts of the multivariate method. The parametric univariate tolerance intervals are calculated as 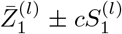, with 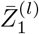 and 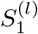 being the location and dispersion parameters of the *l*th ilr variable, *l* = 1, …, *D*, and *c* the tolerance factor. The nonparametric univariate tolerance intervals are determined by taking the union of the middle *k*_0_ equivalent blocks^21^ for each ilr variable. Upon superimposing these tolerance limits on the corresponding uniplots, we may proceed as follows. If the value of the *l*th variable for a composition deemed atypical is not contained in the corresponding interval, then there is indication that the *l*th component may have contributed to the composition’s atypicality.

## 4 Simulation studies

In this section, the coverage performance of the proposed tolerance regions is evaluated through Monte Carlo simulation studies. Since the tolerance region in ℝ^*d*^, where *d* = *D* − 1 given in (14), and the corresponding simplicial tolerance region in 𝒮^*D*^, given in (19), have the same coverage probability, only the region in ℝ^*d*^ is considered.

The simulation study examines the behavior of the classical, robust and *t* tolerance regions under several data-generating settings. First, coverage accuracy is evaluated when the data are generated from a standard multivariate normal distribution and from multivariate *t* distributions with 5 and 10 degrees of freedom. Then, the effect of contaminated training samples is examined by generating the training sample from the contaminated mixture distribution while evaluating the content of the resulting tolerance region with respect to the uncontaminated normal reference distribution. This final setting assesses whether robust estimators lead to tolerance regions that remain close to the intended normal reference region when the estimation sample is contaminated.

The general Monte Carlo procedure for estimating coverage probability is summarized in Algorithm 3.

### Algorithm 3.

**Estimated coverage probability**

For given *d, p, α, n, c*(*d, p, α, n*), large values of *N*_1_ and *N*_2_, a training-sample distribution *G*_train_, and an evaluation distribution *G*_eval_:

1. Generate *N*_1_ independent observations 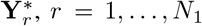, from *G*_eval_. These observations are used to approximate the population content of a constructed tolerance region.
2. Generate a random training sample **Y**_*i*_, *i* = 1, …, *n*, from *G*_train_, and compute the estimates 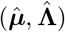 using the method under consideration.
3. Calculate the squared Mahalanobis distances

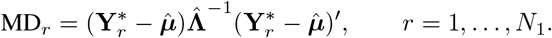
4. Compute the proportion of times MD_*r*_ ≤ *c*(*d, p, α, n*), *r* = 1, …, *N*_1_. Denote this estimated content by 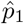.
5. Repeat Steps 2–4 *N*_2_ times to obtain 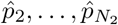.
6. Compute the proportion of times 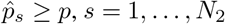. This proportion is the estimated coverage probability, denoted by 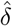.

Algorithm 3 is applied under the following settings. For coverage accuracy under a correctly specified distribution, *G*_train_ = *G*_eval_ = *G*, where *G* is taken to be 𝒩_*d*_(**0, I**), *t*_*d*_(**0, I**, 5), or *t*_*d*_(**0, I**, 10). To study the effect of contaminated training samples on the intended normal reference population, *G*_train_ = *G*_*ϵ*_ where *G*_*ϵ*_ = (1 − *ϵ*) 𝒩_*d*_(**0, I**) + *ϵ 𝒞*_*d*_(**0, I**), itand *G*_eval_ = 𝒩_*d*_(**0, I**). In this setting, the classical and robust tolerance regions are compared to determine whether robust estimation provides improved protection against contamination in the training sample.

In addition to coverage probability, the volumes of the tolerance regions are computed to compare the sizes of the regions produced by different methods. This comparison is useful because two methods may have similar coverage probabilities but produce regions of substantially different sizes. Following Boente and Farall (2008),^22^ the volume is calculated at each Monte Carlo replication and summarized using a relative volume index. For an ellipsoidal tolerance region *T* with dispersion matrix 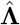, the volume is given by

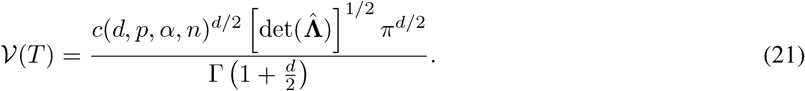

Thus, for each Monte Carlo replication, a volume is obtained for each method. Let

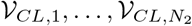

denote the volumes of the classical tolerance regions across the *N*_2_ replications, and let

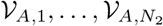

denote the corresponding volumes for another method *A*, where *A* may represent a robust region, a *t*-based region, or a nonparametric region. The relative volume index is defined as

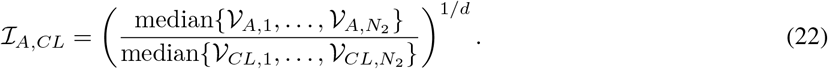

The 1*/d*th root is used to make the index comparable across different dimensions. Values of ℐ_*A,CL*_ greater than one indicate that method *A* produces larger regions than the classical method on average, while values less than one indicate smaller regions.

The simulation settings are as follows. The probability content is fixed at *p* = 0.90, and the confidence level is fixed at 1 − *α* = 0.95. The contamination fraction is set to *ϵ* = 0.05. Dimensions *d* = 2, 3, …, 7 and sample sizes *n* = 50, 100, 200 are considered for the parametric methods. For the computation of coverage probability and volume summaries, *N*_1_ = 10000 and *N*_2_ = 10000 Monte Carlo replications are used.

The simulation results show clear differences among the classical, robust, and *t*-based tolerance regions across the distributional settings. Under the standard multivariate normal distribution, the classical tolerance region performs well because the classical estimators of location and dispersion are well suited to normal data. As shown in Table 1, the estimated coverage probability is close to the nominal confidence level 1 − *α* = 0.95. The robust tolerance regions based on MCD, SD, and S estimators also provide coverage close to the nominal level. The *t*-based regions tend to produce larger regions under normality because they are designed to accommodate heavier-tailed distributions.

**Table 1:**
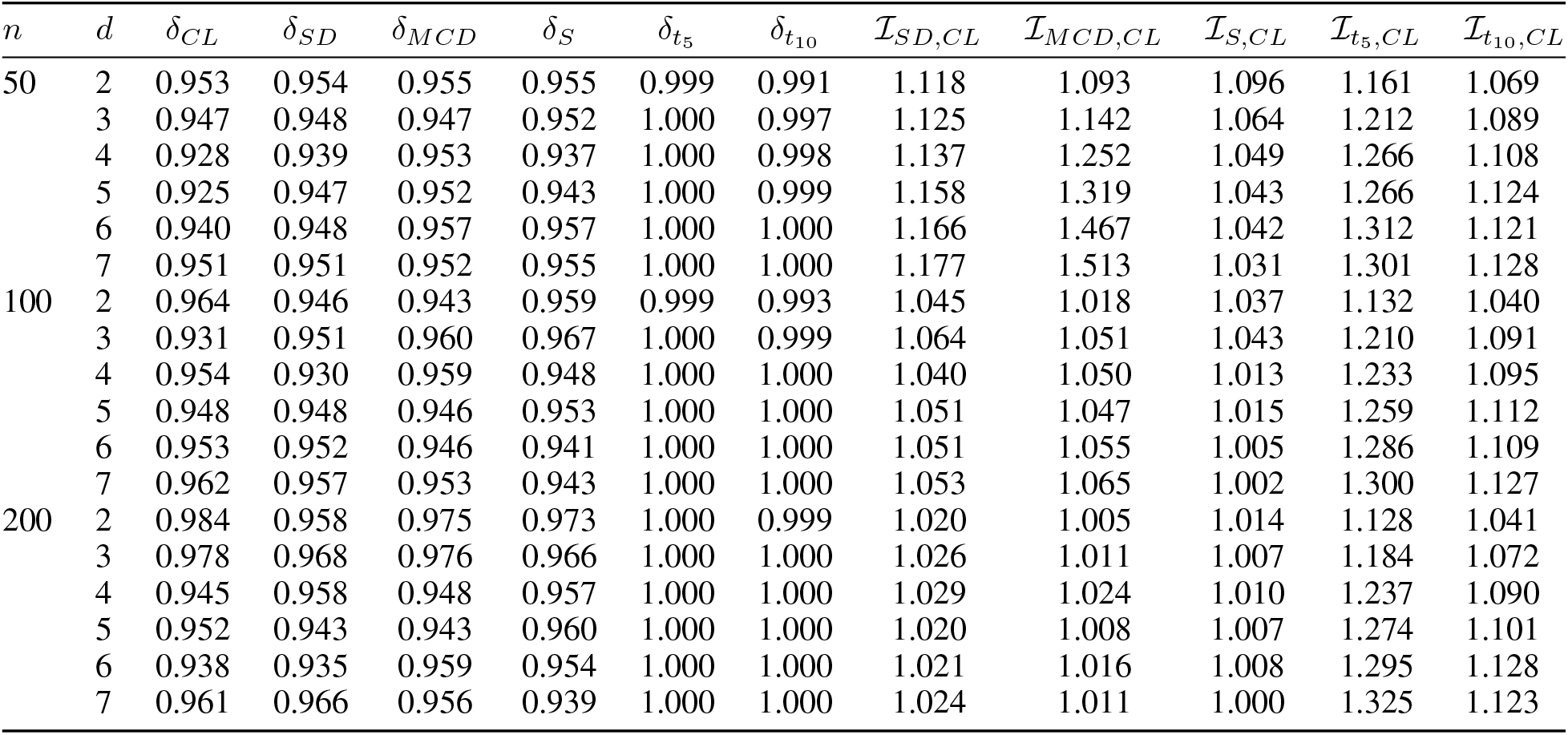
Estimated coverage probabilities of (90, 95)% classical, robust, and *t*-based tolerance regions under 𝒩 _*d*_(0, I).

Under the multivariate *t* distributions, the performance of the classical normal-based tolerance region decreases. As shown in Tables 2 and 3, the heavier tails lead to lower or less stable coverage for the classical and robust region. In contrast, the *t*-based tolerance regions provide improved coverage in these settings, particularly when the degrees of freedom used in the region construction are close to the data-generating distribution.

**Table 2:**
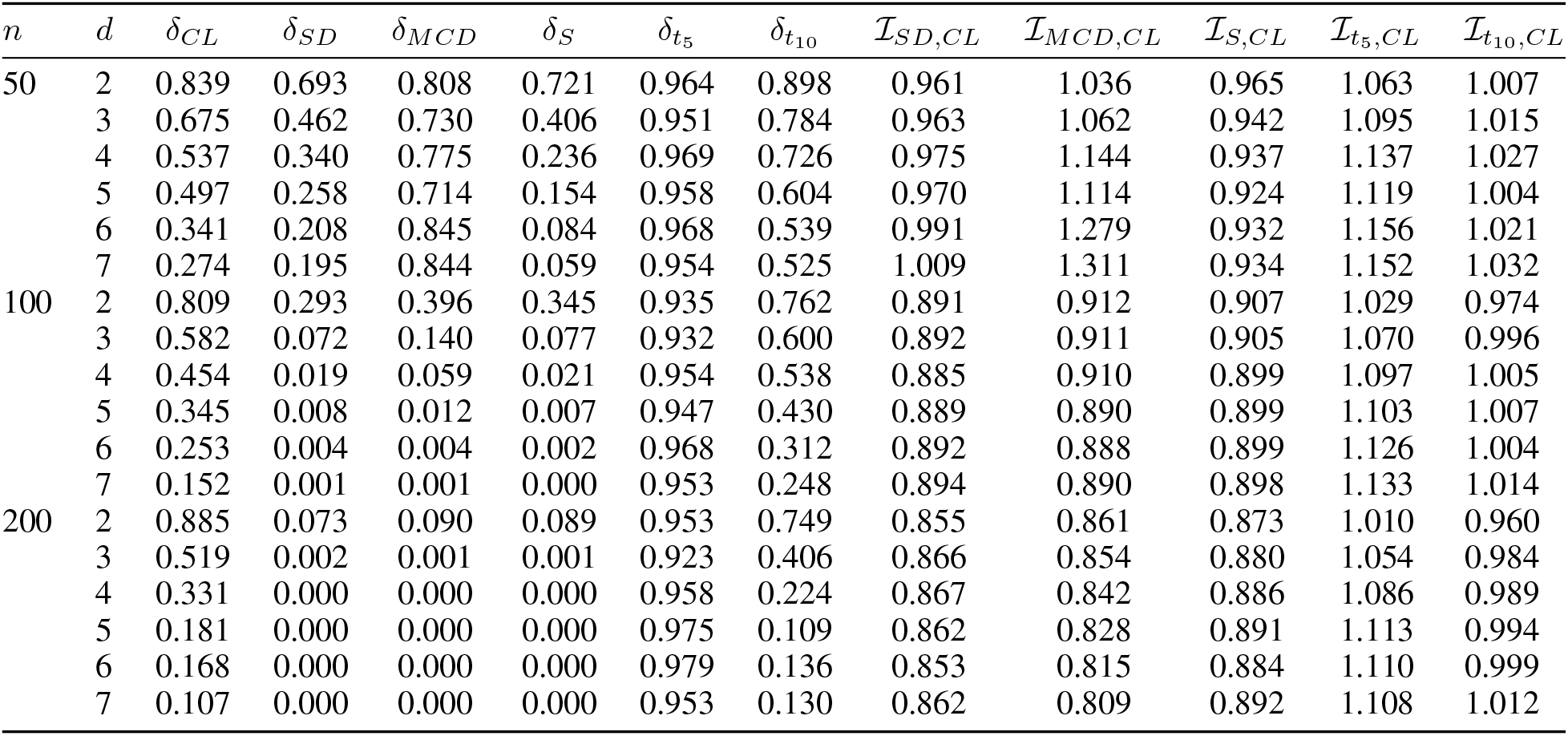
Estimated coverage probabilities of (90, 95)% classical, robust, and *t*-based tolerance regions under *t*_*d*_(0, I, 5)

**Table 3:**
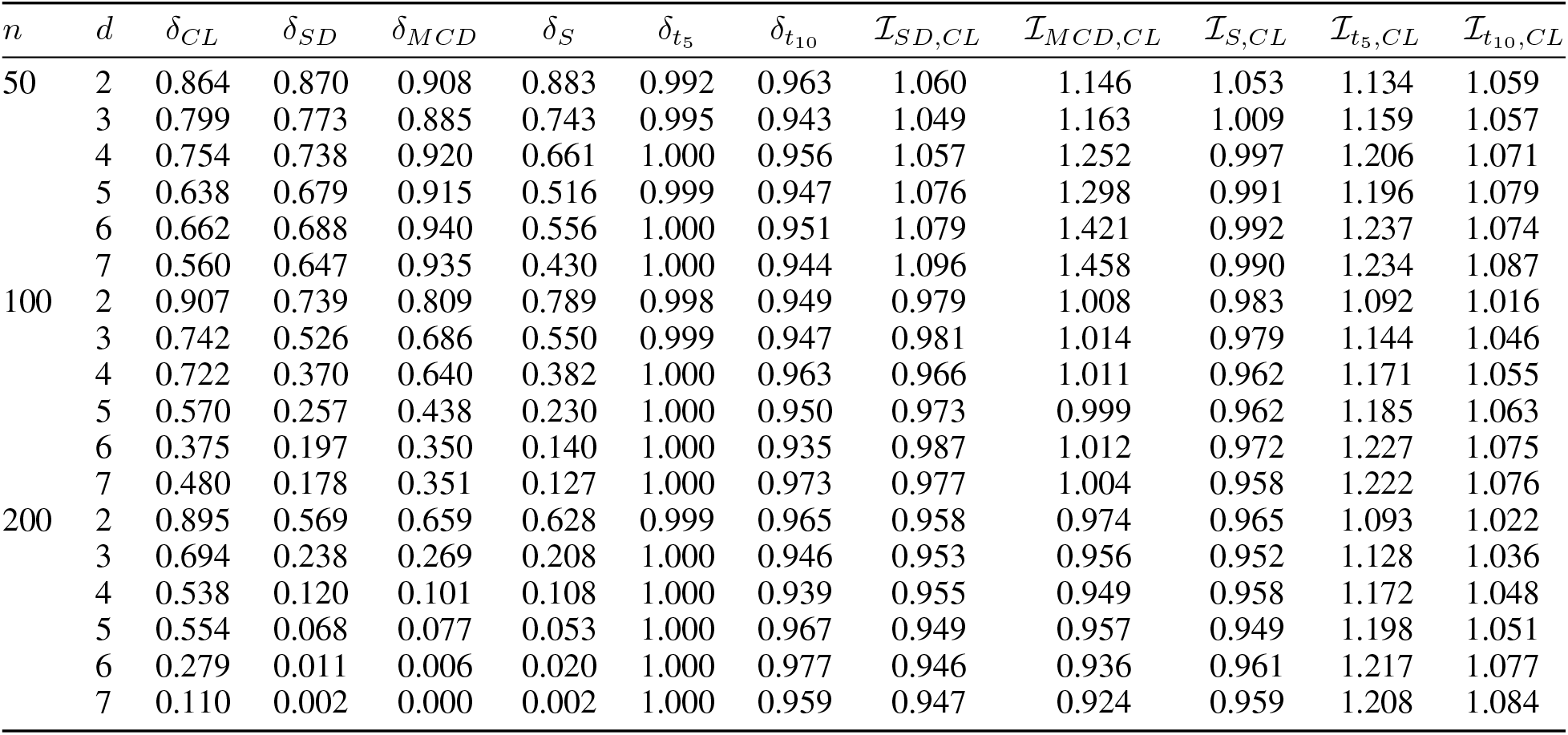
Estimated coverage probabilities of (90, 95)% classical, robust, and *t*-based tolerance regions under *t*_*d*_(0, I, 5)

When the training sample is contaminated but the content is evaluated with respect to the uncontaminated normal reference distribution, the robust regions outperform the classical region. As shown in Table 4, the classical estimates are affected by contamination in the training sample, so the resulting region does not represent the intended normal reference population as well. In contrast, the robust estimators are less affected by contaminated observations, producing tolerance regions that remain closer to the regions obtained from uncontaminated normal data. This setting highlights the practical benefit of robust tolerance regions when constructing reference regions from data that may contain atypical observations.

**Table 4:**
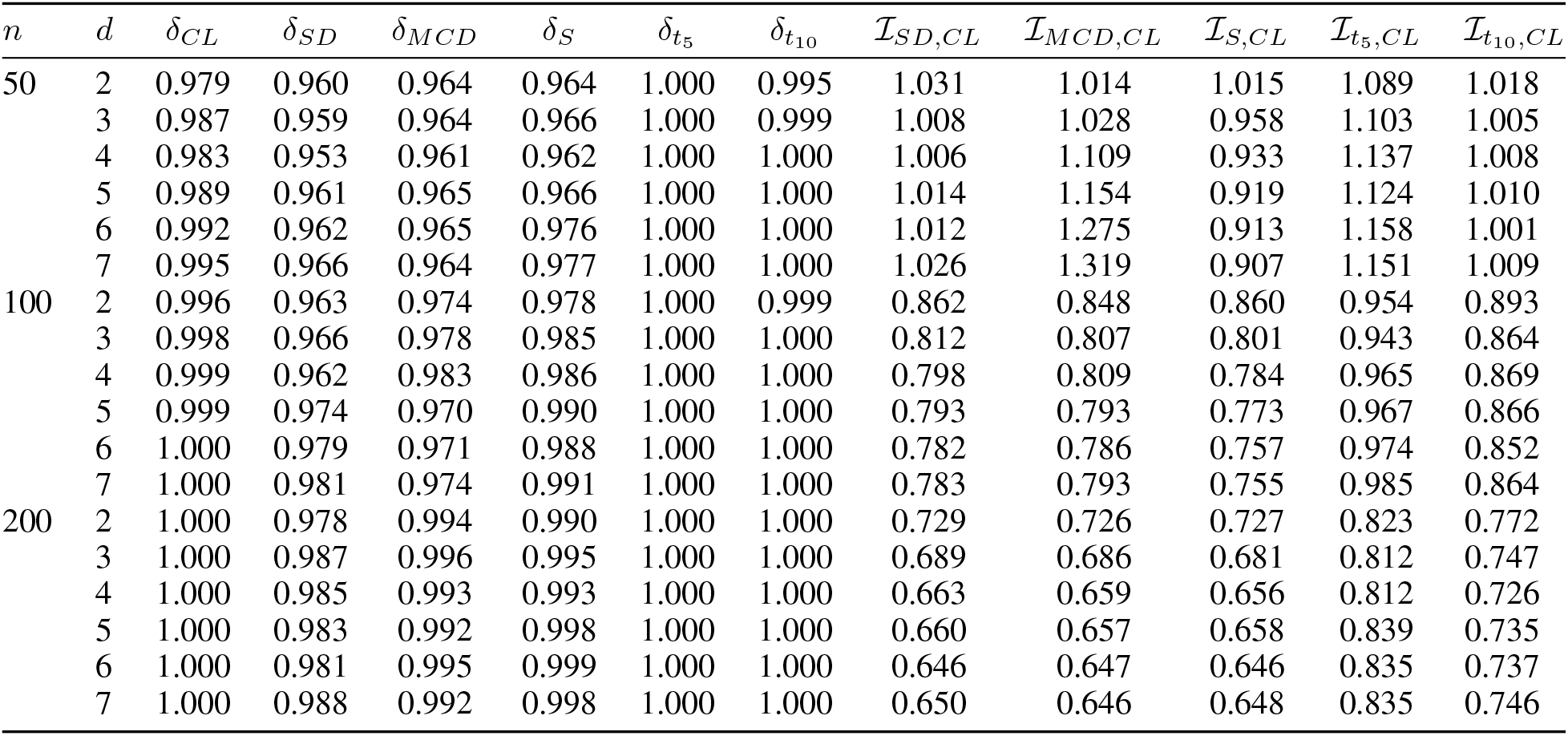
Estimated coverage probabilities of (90, 95)% tolerance regions when the training samples are from (1 − *ϵ*)𝒩_*d*_(0, I) + *ϵ*𝒞_*d*_(0, I) but coverage is evaluated with respect to the 𝒩_*d*_(0, I).

## 5 Reference regions for microbiome data

Microbiome sequencing studies generate multivariate abundance measurements that characterize the microbial composition of individuals in a target population. Such data are typically obtained from biological samples collected at a specific body site, such as the gut, and abundance is quantified at a selected taxonomic rank. Consider microbiome data collected from *n* healthy samples, indexed as *i* = 1, …, *n* and let *D* is the number of taxa of interest for a given taxonomic rank. For the *i*th sample, the observed data consist of a 1 ×*D* vector of nonnegative sequencing counts **W**_*i*_ = [*W*_*i*1_, …, *W*_*iD*_]where *W*_*ij*_ represents the measured abundance of taxon *j*. Because total sequencing depth can vary between samples, direct comparison of raw counts is inappropriate. To obtain scale-invariant measures, the raw counts for each sample are normalized by dividing by their total count, yielding the relative abundance vector **X**_*i*_ = [*X*_*i*1_, …, *X*_*iD*_], where 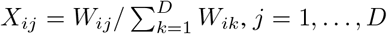. By construction, 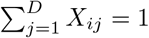, thus **X**_*i*_ lies in the *D*-dimensional simplex 𝒮^*D*^. The methodology developed in Section 3 for constructing tolerance regions for compositional data can be directly applied to these data to construct reference regions for healthy microbiome profiles from the target population. Below we illustrate this application using a gut microbiome dataset. As compositional vectors are constrained to this simplex, the presence of zero components may complicate subsequent log-ratio–based analyses. When zeros occur, established zero-replacement methods can be used to preserve the compositional structure.^39^ The `R` package `compositions`^40^ is used to perform some basic compositional data analysis.

### 5.1 ‘Hitlist’ gut microbiome data

The Hitlist data were downloaded from the study of King et. al,^41^ which established GutFeelingKB, a healthy human gut reference microbiome and abundance profile. The reference catalog contains 157 microorganisms classified into 8 phyla, 18 classes, 23 orders, 38 families, 59 genera, and 109 species. The Hitlist dataset consists of observed taxon counts for *n* = 98 healthy gut samples across these taxa. Of the 98 samples, 50 were obtained from the Human Microbiome Project,^42^ and the remaining samples were collected at George Washington University. For each sample, raw counts were converted to relative abundances by dividing each taxon count by the total count within the sample. Sample SRS016585 was excluded from the analysis due to its extreme compositional structure compared to other samples.

The analysis presented here focuses on the phylum taxonomic rank, which has *D* = 8 distinct taxa: Euryarchaeota, Bacteroidetes, Proteobacteria, Planctomycetes, Verrucomicrobia, Spirochaetes, Actinobacteria, and Firmicutes. Depending on the specific requirements of an application, the approach can be used for any taxonomic rank for which *n >* (*D* − 1). For all analyses, relative abundances were transformed to isometric log-ratio (ilr) coordinates using (2), and (*p*, 1 − *α*) = (0.90, 0.95) was used for constructing tolerance regions. At the phylum level, no zero compositions were observed across samples, and thus zero-replacement was not required prior to log-ratio transformation. To improve the clarity in visualization, samples were relabeled according to their corresponding observation numbers. A complete mapping of the original sample identifiers to the assigned observation numbers is provided Supplementary Materials.

### 5.2 Reference regions for a three-part subcomposition

We first consider a simpler illustration with D=3, for which the compositions in the simplex and their two ilr coordinates can be easily visualized. The chosen Three-part subcomposition consists of Bacteroidetes (*X*_1_), Firmicutes (*X*_2_), and Proteobacteria (*X*_3_).

Figure 1a displays the observed data in a ternary diagram. Bacteroidetes and Firmicutes tend to dominate the composition and Bacteroidetes tend to have more weight than Firmicutes. Moreover, the variation in the ratio of Bacteroidetes and Proteobacteria is smaller than the other two ratios. These observations are confirmed by the geometric center and dispersion matrix of the data. see Appendix 3 of the Supplementary Document for their definitions:

**Figure 1:**
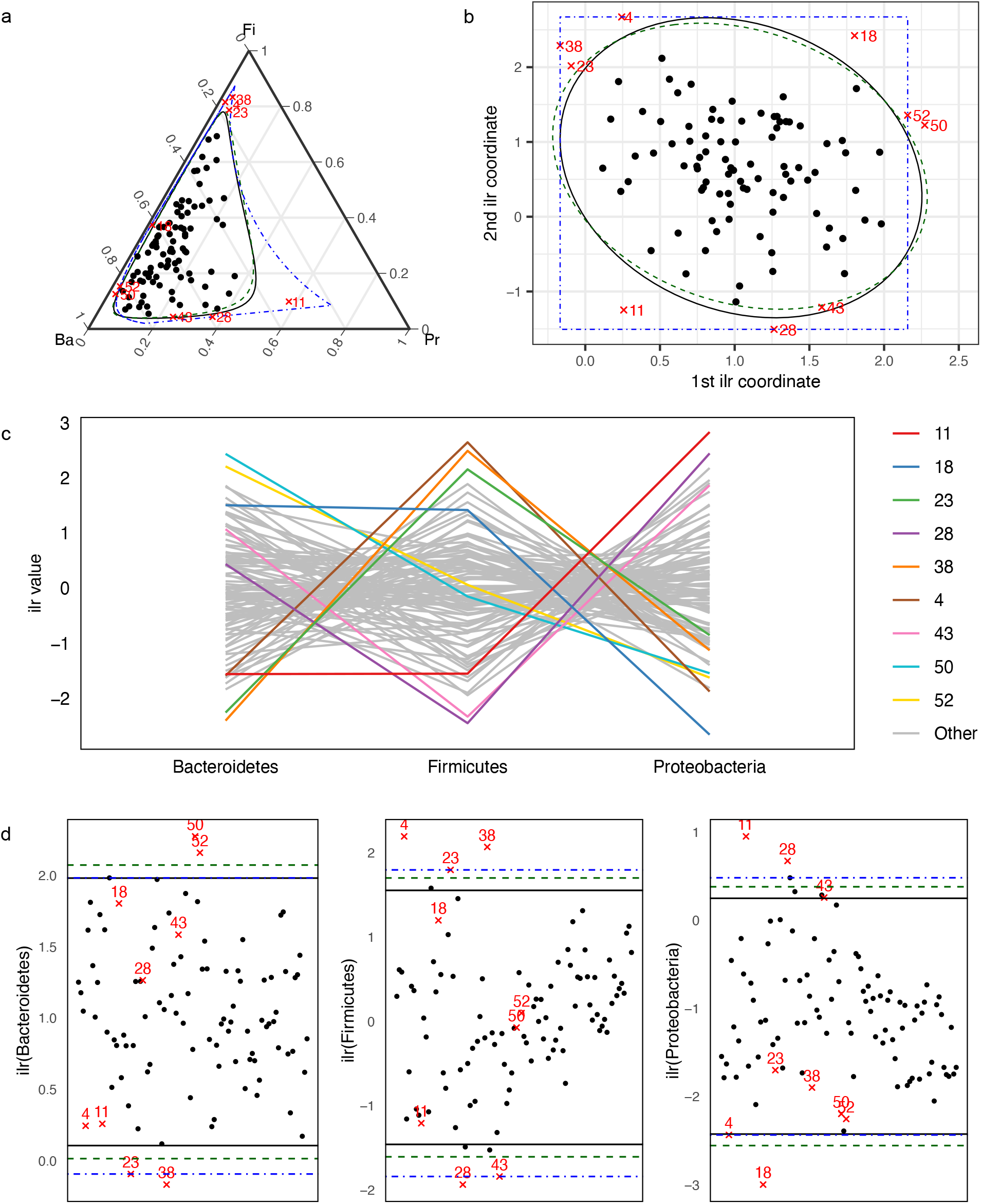
Atypical observations for the 3-part subcomposition—Bacteroidetes (Br), Firmicutes (Fi), and Proteobacteria (Pr)—and the corresponding reference regions. (90,95)% parametric reference regions based on the classical approach (black solid line) and the robust approach using the SD estimator (green dashed line), and nonparametric reference regions (blue dot–dash line). Observations falling outside the reference regions are marked in red. **a** Ternary diagram for original data in 𝒮^3^ and the simplicial reference region. **b** Scatter plot of ilr coordinates in ℝ^2^ and the ellipsoidal reference region. **c** Parallel plot of ilr variables, with observations inside the reference region shown in light gray. **d** Uniplots of ilr variables with superimposed univariate reference intervals.

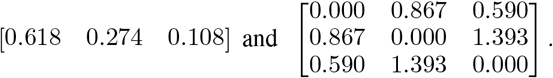

Also, none of the off-diagonal elements in the matrix are particularly small, indicating that there is no evidence of strong codependence between any taxa pair.

Figure 1b displays a scatter plot of the two ilr coordinates. The first coordinate (*Z*_1_) reflects the difference in abundance of Bacteroidetes compared to Firmicutes and Proteobacteria. Whereas, the second coordinate (*Z*_2_) reflects the difference in abundance of Firmicutes compared to Proteobacteria. Most of the values for the two coordinates are positive, indicating that Bacteroidetes tend to be more abundant than the other two taxa and Firmicutes are more abundant than Proteobacteria, which is consistent with the above finding. The normality assumption was evaluated using the ilr coordinates before constructing the tolerance region. As reported in Appendix 4 of the Supplementary Document, the assumption of normality appears reasonable. Classical, robust, and nonparametric tolerance regions were constructed to allow comparison of the three approaches. For the robust method, the Stahel–Donoho (SD) estimator was used due to its superior performance in the simulation studies. The maximum likelihood estimators for both classical and robust methods are as follows:

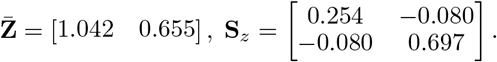

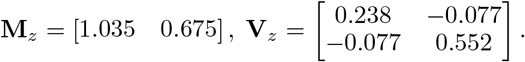

Figure 1b also shows the ellipsoidal, nonparametric and simplicial reference regions with *p* = 0.90 and *α* = 0.95, as given by (14), (17), and (19) respectively. These tolerance regions are constructed to contain at least 90% of the healthy population’s gut subcompositions with 95% confidence. Consequently, approximately 10% of compositions from healthy individuals are expected to fall outside the reference regions, consistent with the specified population coverage. The tolerance factors *c*(2, 0.90, 0.05, 97) for the classical and robust regions given by (6) and (10), are 5.781 and 6.618, respectively. The nonparametric reference region is a union of 93 equivalent blocks and the exact confidence level is 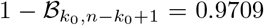. The reference limits for the 1st and 2nd ilr coordinates are (−0.168, 2.158) and (−1.506, 2.672), respectively.

Figure 1 and Table 5 summarize the observations that fall outside the reference regions. Table 5 reports their squared Mahalanobis distances from the center, the corresponding *p*-indices defined in (4), and the number of equivalent blocks required for the nonparametric region. Eight observations: #4, 11, 18, 23, 28, 38, 50, and 52 fall outside the classical reference region. One additional observation (#43) falls outside the robust reference region, but it is very close to the boundary. As expected, for these observations the squared Mahalanobis distances exceed the tolerance factor values, and the corresponding *p*-indices exceed the probability content of 0.90. To interpret a *p*-index, consider #4 whose *p*-index is 0.943 for classical region. This observation falls among the most extreme 6% of the healthy compositions. The *p*-indices for other observations can be interpreted similarly. Observation #11 stands out as the most atypical point, while #43 is the least atypical. Observation #50 is the only observation that falls outside the nonparametric region and #52, 4, 38 and 28 are on the boundary. The center values are close across all three regions but the nonparametric region is clearly bigger than two parametric regions, resulting in fewer atypical observations. However, since the normality assumption holds for the data, the parametric regions are preferable and the two versions of it differ little.

**Table 5:** Observations that fall outside the (90, 95)% classical, robust, and nonparametric reference regions for 3-part subcomposition.

| ID | Sample ID | Classical Region |  | Robust Region |  | Nonparametric Region |  |
| --- | --- | --- | --- | --- | --- | --- | --- |
|  |  | MD<sup>2</sup> | *p*-index | MD<sup>2</sup> | *p*-index | *k*<sub>0</sub> | *p*-index |
| 4 | SRS011302 | 7.156 | 0.943 | 8.373 | 0.948 |  |  |
| 11 | SRS013215 | 9.312 | 0.977 | 11.497 | 0.979 |  |  |
| 18 | SRS014313 | 8.277 | 0.963 | 10.015 | 0.968 |  |  |
| 23 | SRS014979 | 6.596 | 0.932 | 7.178 | 0.919 |  |  |
| 28 | SRS015369 | 6.710 | 0.931 | 8.632 | 0.947 |  |  |
| 38 | SRS016495 | 8.114 | 0.959 | 8.936 | 0.956 |  |  |
| 43 | SRS017247 |  |  | 6.797 | 0.906 |  |  |
| 50 | HMB_01-1 | 7.327 | 0.949 | 8.149 | 0.937 | 93 | 0.922 |
| 52 | HMB_01-3 | 6.558 | 0.930 | 7.373 | 0.921 |  |  |

Because we can plot the original data in a ternary diagram and superimpose the reference region, gaining insight into the atypicality of the points outside the region is straightforward. As shown in Figure 1a, the simplicial region shows why the nine compositions are flagged as atypical. Specifically, #11 is unusually high and #18 is nearly zero in Proteobacteria; #4, 23, and 38 are unusually high in Firmicutes, while #28 and #43 are low in Firmicutes; #50 and #52 are high in Bacteroidetes, whereas #4, 23, and 38 are low in Bacteroidetes. It is now of interest to use graphical and numerical summaries of the ilr variables, as described in Section 3.3, for the same purpose. The parallel plot of the three ilr variables in Figure 1c displays the multivariate structure of the data. Observations inside the reference region share a common pattern, whereas those outside differ substantially. The results are consistent with the ternary diagram. Observations with high compositions of Bacteroidetes, Proteobacteria, or Firmicutes correspond to high ilr values in the parallel plot, whereas those with low compositions correspond to low ilr values.

Figure 1d displays uniplots of the ilr variables with the corresponding univariate (90, 95)% classical, robust, and nonparametric reference limits superimposed. The uniplots are constructed as described in Section 3.3 to identify which components contribute to atypicality. An observation falling outside the reference intervals is considered atypical due to an unusual composition in that specific component. For Bacteroidetes, #50 and #52 lie above the upper limits of all three reference intervals, indicating unusually high proportions of this taxon, whereas #38 falls below the lower limit, indicating an unusually low proportion. In Firmicutes, #4 and #38 exceed the upper limit of all three univariate reference intervals, indicating elevated compositions, while #28 falls below the lower limit, suggesting a comparatively low proportion. In Proteobacteria, #11 and #28 lie above the upper limit whereas #18 falls below the lower limit.

Some observations, such as #23 and #38, lie on the boundary of the nonparametric reference intervals. However, the nonparametric limits are wider than the parametric ones, and, as discussed earlier, the parametric intervals are preferred since the normality assumption is reasonable.

Taken together, we see that the conclusions based on ilr variables for *D* = 3 are in line with those based on the ternary diagram. This confirms the utility of the ilr variables in gaining insights into atypicality, especially for *D >* 3 in which case visualizing data in the simplex is difficult.

### 5.3 Reference regions for the full eight-part composition

In this section, we constructed reference regions for the full composition with *D* = 8 parts. The figures presented in Appendix 4 of the Supplementary Document suggests that the marginal distributions are reasonably consistent with normality. However, the chi-squared Q–Q plot for normal distribution indicates the presence of heavier tails, implying that the joint distribution may deviate from multivariate normality. To account for the heavier tails, we modeled the data using a *t* distribution with 10 degrees of freedom.

Estimates for the eight-part compositions are presented in Appendix 4 of the Supplementary Document. The compositions are dominated by Bacteroidetes, with Euryarchaeota, Planctomycetes, and Spirochaetes representing smaller proportions. Moreover, the variation in the ratio of Proteobacteria to Planctomycetes is the highest, followed by the variation between Planctomycetes and Verrucomicrobia. The variation between Bacteroidetes and Actinobacteria is relatively small.

Classical and *t*-reference regions were constructed with *p* = 0.90 and *α* = 0.95 for the 8-part composition. Based on the goodness-of-fit considerations, the *t* reference region is selected as the primary reference region. The classical reference region is included only to facilitate comparison with the *t* reference regions. The tolerance factors *c*(7, 0.90, 0.05, 97) for the classical and *t* regions given by (6) and (14), are 14.899 and 16.952, respectively. Nonparametric regions are not reported because they do not exist for the current sample configuration (*n, D* − 1) = (97, 7). When *D* = 8, a minimum sample size of *n* ≥ 227 is required for the nonparametric region to exist.

Because of the high dimensionality, the reference regions cannot be visualized using ternary diagrams. However, it is straightforward to determine whether a given composition lies inside or outside the reference region, and to assess its distance from the center, using the *p*-index. Out of the 97 observations, 6 observations (#9, 11, 18, 23, 36, and 96) fall outside the *t* reference region. In addition, observations #10 and #38 fall outside the normal reference region, resulting in a total of 8 observations outside the normal region. The Mahalanobis distances from the center and the corresponding *p*-indices are reported in Table 6. For these observations, the distances exceed the corresponding tolerance factor values, and the *p*-indices are greater than the 0.90 probability content threshold. The *p*-indices for observations #10 and #38 indicate that they lie close to the boundary of the normal reference region, whereas observation #36 has a *p*-index close to one, suggesting extreme atypicality.

**Table 6:**
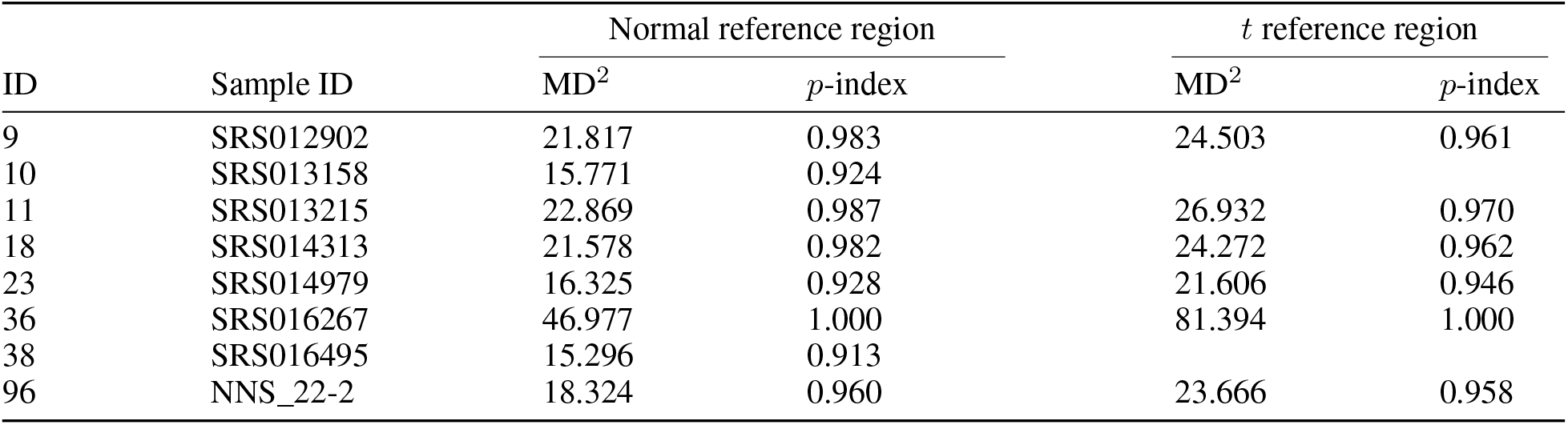
Observations that fall outside the (90, 95)% normal and *t* reference regions for 8-part composition.

Figure 2a displays the parallel plot for the eight ilr variables. The multivariate structure of the atypical observations differs from that of the remaining observations. The uniplots in Figure 2b, together with Table 7, identify observations that fall outside the univariate normal and *t* reference intervals and help determine which variables contribute to these deviations.

**Figure 2:**
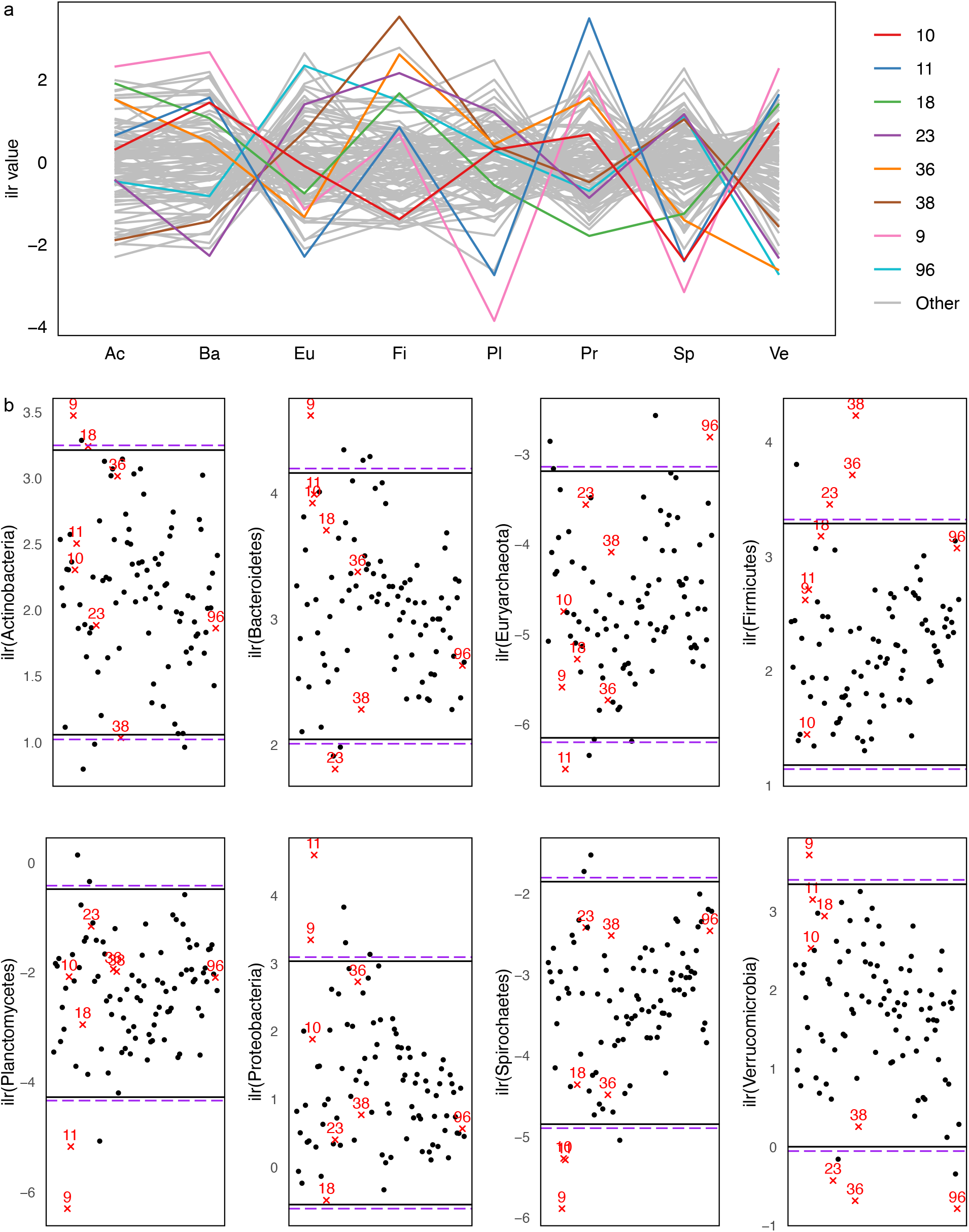
Atypical observations for the 8-part composition with (90,95)% classical reference regions (black solid line) and *t* reference regions (purple long–dash line). Observations falling outside the reference regions are marked in red. **a** Parallel plot of ilr variables, with observations inside the reference region shown in light gray. **b** Uniplots of ilr variables with superimposed univariate reference intervals.

**Table 7:** Uniplot location of observations that fall outside univariate normal and *t* reference intervals for 8-part composition. H indicates an observation above the upper limit, L indicates an observation below the lower limit, and an empty cell indicates an observation within the limits.

| | Normal reference region | | | | | | | | $t$ reference region | | | | | |
| --- | --- | --- | --- | --- | --- | --- | --- | --- | --- | --- | --- | --- | --- | --- |
|  | 9 | 10 | 11 | 18 | 23 | 36 | 38 | 96 | 9 | 11 | 18 | 23 | 36 | 96 |
| Actinobacteria | H |  |  | H |  |  | L |  | H |  |  |  |  |  |
| Bacteroidetes | H |  |  |  | L |  |  |  | H |  |  | L |  |  |
| Euryarchaeota |  |  | L |  |  |  |  | H |  | L |  |  |  | H |
| Firmicutes |  |  |  |  | H | H | H |  |  |  |  | H | H |  |
| Planctomycetes | L |  | L |  |  |  |  |  | L | L |  |  |  |  |
| Proteobacteria | H |  | H |  |  |  |  |  | H | H |  |  |  |  |
| Spirochaetes | L | L | L |  |  |  |  |  | L | L |  |  |  |  |
| Verrucomicrobia | H |  |  |  | L | L |  | L | H |  |  | L | L | L |

Observation #9 appears atypical due to high Actinobacteria, Bacteroidetes, Proteobacteria, and Verrucomicrobia together with very low Planctomycetes and Spirochaetes. Observation #11 shows high Proteobacteria and low Euryarchaeota, Planctomycetes, and Spirochaetes, while observation #18 has low Proteobacteria. Observation #23 is characterized by high Firmicutes and low Bacteroidetes and Verrucomicrobia, and observation #36 shows high Firmicutes with low Verrucomicrobia. Finally, observation #96 has high Euryarchaeota and low Verrucomicrobia.

### 5.4 Validation of reference regions in health vs. disease

We applied the proposed reference region methodology to two microbiome datasets at the phylum level to compare the microbiome profiles of healthy individuals with those of patients. Specifically, we constructed reference regions using samples from healthy individuals and then assessed whether patient samples could be identified as atypical. We also explored which taxa might be contributing to these deviations.

Our analysis focused on subcompositions formed from the most abundant phyla and taxa that showed differential abundance between healthy and diseased individuals. To identify suitable subcompositions, we first selected taxa with fewer than 20% zero values across all samples and then constructed reference regions for all possible combinations of these taxa. Among these, we chose the subcomposition that classified the highest proportion of diseased observations as atypical. This approach allowed us to identify subcompositions that best differentiated healthy and diseased groups. We then examined centered log-ratio (CLR)-transformed boxplots presented in Appendix 4 of the Supplementary Document to visually assess distributional differences between the two groups for these subcompositions. In addition, we reviewed the existing literature to determine whether the taxa in the selected subcomposition had previously been reported to be associated with the diseases under study.

For each dataset, we first constructed the normal, robust, and *t* reference regions with *p* = 0.9 and *α* = 0.05 using data from healthy individuals, following the method outlined in Section 3. Next, we assessed how many patient samples fall outside these reference regions by computing the Mahalanobis distance from the center of the reference region for each observation and calculating the corresponding *p*-index. Finally, we examined the ilr variables to identify potential taxa that may explain why these patient samples appear atypical. This method is particularly useful when the selected subcomposition contains taxa that show differential abundance between healthy and diseased groups.

#### 5.4.1 Liver cirrhosis study

The first dataset is from the study conducted by Qin et al.^43^ and focuses on gut microbiome profiles in patients with liver cirrhosis. The dataset includes samples from 123 Chinese patients diagnosed with liver cirrhosis alongside 114 healthy control individuals. Bacteroidetes, Firmicutes, and Proteobacteria stand out as the most differentially abundant taxa and we carried out the analysis using this subcomposition. Literature suggest that there are significant differences in the relative abundances of these taxa between patients with cirrhosis and healthy individuals.^44,45^

The robust reference region identified the highest number of atypical individuals, flagging 41 patients as atypical observations. Figure 3a, 3b displays the observed data along with the constructed robust reference region for liver cirrhosis. The figure clearly shows differences in the microbial compositions between healthy individuals and patients. For healthy individuals, Bacteroidetes tend to be more abundant than Firmicutes and Proteobacteria, while in patients, the abundance of Bacteroidetes is lower compared to healthy individuals. Additionally, some patients exhibit significantly higher levels of Proteobacteria and Firmicutes, which differentiate them from healthy individuals. Patients that fall outside the robust reference regions, their squared Mahalanobis distances (MD^2^) from the center and their *p*-indices are presented in Table 8. Very high *p*-indexs supports their classification as atypical due to their significant deviation from the reference region.

**Figure 3:**
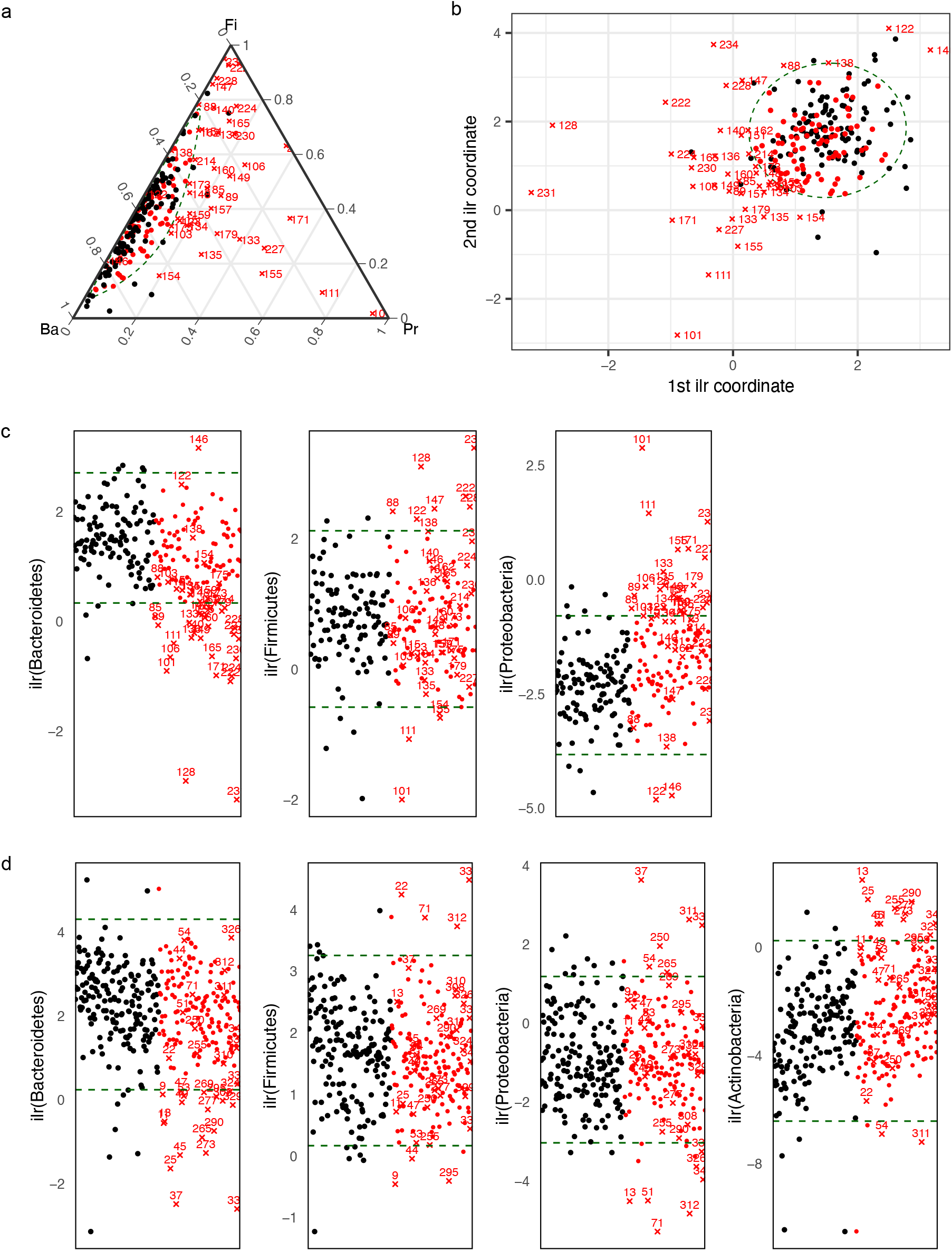
Atypical observations for the 3-part subcomposition, along with the corresponding (90, 95)% parametric reference region obtained using the robust SD-based approach for patient data. Diseased observations are shown as red dots, and diseased atypical observations are shown as red crosses. **a** Ternary diagram of the original data in 𝒮^3^ with the simplicial reference region for the liver cirrhosis study. **b** Scatter plot of the ilr coordinates in ℝ^2^ with the ellipsoidal reference region for the liver cirrhosis study. **c** Uniplots of the ilr variables with superimposed univariate robust reference intervals. for the liver cirrhosis study. **d** Uniplots of the ilr variables with superimposed univariate robust reference intervals for the type 2 diabetes study.

**Table 8:** Observations that fall outside the (90, 95)% robust reference regions for the liver cirrhosis study and their uniplot locations. H indicates an observation above the upper limit, L indicates an observation below the lower limit, and an empty cell indicates an observation within the limits.

| ID | MD<sup>2</sup> | *p*– index | Ba | Fi | Pr |
| --- | --- | --- | --- | --- | --- |
| 85 | 11.1227 | 0.9902 | L |  | H |
| 88 | 8.0150 | 0.9636 |  | H |  |
| 89 | 14.4614 | 0.9967 | L |  | H |
| 101 | 78.5896 | 1.0000 | L | L | H |
| 103 | 6.9416 | 0.9438 |  |  |  |
| 106 | 22.1604 | 0.9998 | L |  | H |
| 111 | 41.9698 | 1.0000 | L | L | H |
| 115 | 6.8084 | 0.9369 |  |  |  |
| 122 | 17.7374 | 0.9992 |  | H | L |
| 123 | 7.3539 | 0.9485 |  |  | H |
| 128 | 76.8469 | 1.0000 | L | H | H |
| 133 | 19.4517 | 0.9998 | L |  | H |
| 134 | 9.0871 | 0.9753 |  |  | H |
| 135 | 14.0747 | 0.9972 |  |  | H |
| 136 | 13.5726 | 0.9959 | L |  |  |
| 138 | 6.3201 | 0.9285 |  |  |  |
| 140 | 11.6455 | 0.9898 | L |  |  |
| 146 | 19.0174 | 0.9992 | H |  | L |
| 147 | 11.1703 | 0.9897 | L | H |  |
| 148 | 7.4370 | 0.9538 |  |  |  |
| 149 | 16.6927 | 0.9988 | L |  | H |
| 151 | 7.5431 | 0.9556 | L |  |  |
| 154 | 11.0624 | 0.9888 |  | L | H |
| 155 | 25.9822 | 0.9999 | L | L | H |
| 157 | 12.6844 | 0.9942 | L |  | H |
| 159 | 8.7311 | 0.9737 |  |  | H |
| 160 | 12.3854 | 0.9936 | L |  | H |
| 162 | 6.4038 | 0.9266 | L |  |  |
| 165 | 18.8627 | 0.9994 | L |  | H |
| 171 | 34.5759 | 1.0000 | L |  | H |
| 173 | 6.8640 | 0.9415 |  |  |  |
| 175 | 6.3235 | 0.9276 |  |  |  |
| 179 | 15.0167 | 0.9973 | L |  | H |
| 214 | 6.9132 | 0.9377 | L |  |  |
| 222 | 28.0884 | 1.0000 | L | H |  |
| 224 | 25.2427 | 0.9999 | L |  | H |
| 227 | 24.7605 | 0.9999 | L |  |  |
| 228 | 13.5679 | 0.9955 | L | H |  |
| 230 | 20.3026 | 0.9997 | L |  | H |
| 231 | 92.9050 | 1.0000 | L |  | H |
| 234 | 24.1460 | 0.9999 | L | H |  |

Figure 3c displays uniplots of the ilr variables, highlighting which variables may have caused the patients to lie outside the multivariate tolerance region. As shown in uniplots and Table 8, many patients exhibit lower levels of Bacteroidetes and higher levels of Proteobacteria compared to healthy individuals. These observation consistent with previous studies on microbial profiles in liver cirrhosis.^44,45^ Firmicutes levels, on the other hand, vary among patients—some have higher levels than healthy individuals, while others have lower levels.

#### 5.4.2 Type 2 diabetes study

The second dataset focuses on the gut microbiome composition in patients with type 2 diabetes, as detailed in the study by Qin et al.^46^ This dataset consist of gutmicrobiome profiles for 170 patients diagnosed with type 2 diabetes and 174 healthy control individuals from southern China. We constructed the reference regions for a subcomposition of four most abundant Phyla—Bacteroidetes, Firmicutes, Proteobacteria, and Actinobacteria. Previous studies have shown significant differences in the relative abundances of these phyla between individuals with type 2 diabetes and healthy controls.^47^ The robust reference region identifies the 33 patients as atypical.

Table 9 presents observations that fall outside the robust reference regions, along with their squared Mahalanobis distances (MD^2^) from the center, *p*-indices, and the ilr variables contributing to their atypicality. Very high *p*-values is a indication that these atypical observations are far from the constructed reference regions. As shown in Table 9 and Figure 3d, patients exhibit lower abundance of Bacteroidetes and higher abandance of Actenobacteria compared to healthy individuals.^48^ Additionally, the abundance of Proteobacteria and Firmicutes differs between healthy individuals and Type 2 diabetes patients, with variations observed as either higher or lower.

**Table 9:** Observations that fall outside the (90, 95)% robust reference regions for the type 2 diabetes study and their uniplot locations. H indicates an observation above the upper limit, L indicates an observation below the lower limit, and an empty cell indicates an observation within the limits.

| ID | $MD^2$ | $p$-index | Ba | Fi | Pr | Ac |
| --- | --- | --- | --- | --- | --- | --- |
| 9 | 13.293 | 0.991 | L | L |  |  |
| 11 | 14.962 | 0.995 | L |  |  |  |
| 13 | 23.895 | 1.000 | L |  | L | H |
| 22 | 29.923 | 1.000 |  | H |  |  |
| 25 | 25.670 | 1.000 | L |  |  | H |
| 37 | 100.817 | 1.000 | L |  | H |  |
| 44 | 8.709 | 0.942 |  | L |  |  |
| 45 | 24.121 | 1.000 | L |  |  | H |
| 47 | 9.903 | 0.964 | L |  |  |  |
| 49 | 9.637 | 0.962 | L |  |  |  |
| 51 | 13.480 | 0.992 |  |  | L | H |
| 53 | 10.371 | 0.968 | L |  |  |  |
| 54 | 8.946 | 0.949 |  |  | H | L |
| 71 | 19.007 | 0.999 |  | H | L |  |
| 250 | 10.563 | 0.972 |  |  | H |  |
| 255 | 9.868 | 0.962 |  |  |  | H |
| 265 | 26.483 | 1.000 | L |  | H |  |
| 269 | 19.678 | 0.999 | L |  |  |  |
| 273 | 21.758 | 1.000 | L |  |  | H |
| 277 | 11.061 | 0.976 | L |  |  | H |
| 290 | 18.461 | 0.999 | L |  |  | H |
| 295 | 12.668 | 0.988 | L | L |  |  |
| 308 | 13.758 | 0.992 | L |  |  |  |
| 310 | 9.157 | 0.953 |  |  |  |  |
| 311 | 17.745 | 0.999 |  |  | H | L |
| 312 | 15.863 | 0.997 |  | H | L |  |
| 324 | 9.068 | 0.950 | L |  |  |  |
| 326 | 10.610 | 0.973 |  |  | L |  |
| 329 | 9.745 | 0.961 | L |  |  | H |
| 337 | 19.389 | 0.999 |  | H | L |  |
| 338 | 75.325 | 1.000 | L |  | H |  |
| 339 | 7.960 | 0.924 |  |  |  |  |
| 340 | 9.716 | 0.962 |  |  | L | H |

## 6 Discussion

This study introduces a methodology for constructing tolerance regions for compositional data, addressing the need for reference regions in microbiome research. These reference regions serve as benchmarks for identifying deviations from normal microbiome composition, detecting unusual profiles, and supporting the discovery of disease-associated microbiome patterns. The proposed framework directly accounts for the compositional nature of microbiome data. Unlike methods that focus only on uncertainty in estimated parameters, the proposed framework constructs regions designed to contain a specified proportion of the healthy population with a given level of confidence. Moreover, despite the high dimensionality of microbiome data, the approach enables clear interpretation of atypicality by introducing a p-index, which quantifies how far a profile lies from the reference region. Using ilr-transformed variables, graphical displays, and univariate confidence intervals, the methodology also helps identify which components contribute most to these deviations.

The proposed framework incorporates both parametric and nonparametric approaches for constructing tolerance regions. Parametric methods are effective when ilr-transformed data follow an approximate elliptical distribution, and they can yield smaller regions while maintaining the desired coverage. In cases where microbiome data contain outliers or contaminated observations, robust estimators of location and dispersion can be incorporated as an alternative to classical estimators, allowing the resulting reference regions to better reflect the target healthy population. Extending the framework beyond the normal distribution to more general elliptical distributions also provides additional flexibility. For example, the simulation results show that tolerance regions based on the multivariate t distribution outperform normal-based regions when the data exhibit heavier tails. However, parametric methods can be sensitive to deviations from the assumed distribution. In such cases, nonparametric approaches provide a flexible alternative by avoiding distributional assumptions, although they may require larger sample sizes and can become less efficient in higher-dimensional settings.

The performance of the method also depends on the selected subcomposition of taxa. If the chosen taxa do not adequately capture differences between healthy and patient groups, the ability to identify atypical profiles may be limited. This underscores the importance of identifying discriminative taxa. One possible approach is to use logistic regression to construct a score that separates healthy and patient profiles, followed by the construction of a univariate tolerance interval based on healthy samples. In addition, microbiome composition is influenced by a range of covariates, including diet, lifestyle, medication use, and environmental exposures. Extending the framework to incorporate covariates through a regression-based tolerance-region approach would allow for more targeted and context-specific reference regions.

Overall, the proposed framework demonstrates that tolerance regions for compositional data offer a statistically meaningful and interpretable approach for defining healthy microbiome reference regions.

