## Supplemental Documents for "Tolerance Regions for Compositional Data with Application to Reference Regions for Healthy Microbiome Profiles"

April 26, 2026

#### Contents

Appendix 1: Additional logratio transformations.

Appendix 2: Proofs of propositions.

Appendix 3: Some basic compositional data analysis concepts

Appendix 4: Additional results for reference regions for microbiome data

### Appendix 1: Additional logratio transformations

This section briefly reviews the centered logratio (clr) and additive logratio (alr) transformations. Let $\mathcal{S}^D$ denote the $D$-part simplex, and let $\mathbf{x} = [x_1, \ldots, x_D] \in \mathcal{S}^D$ be a composition. The simplex $\mathcal{S}^D$ is a vector space with operations of *perturbation* and *powering* serving as vector addition and scalar multiplication, respectively. Perturbation of $\mathbf{x} \in \mathcal{S}^D$ by $\mathbf{y} \in \mathcal{S}^D$ is defined as

$$\mathbf{x} \oplus \mathbf{y} = C[x_1 y_1, \ldots, x_D y_D].$$

Powering of $\mathbf{x} \in \mathcal{S}^D$ by a constant $a \in \mathbb{R}$ is defined as

$$a \odot \mathbf{x} = C[x_1^a, \ldots, x_D^a].$$

The vector space is equipped with Aitchison inner product and the associated norm and distance, making it a Euclidean space. For $\mathbf{x}, \mathbf{y} \in \mathcal{S}^D$, the three quantities are respectively defined as follows:

$$\langle \mathbf{x}, \mathbf{y} \rangle_{\mathrm{a}} = \frac{1}{2D} \sum_{j=1}^{D} \sum_{k=1}^{D} \ln \frac{x_j}{x_k} \ln \frac{y_j}{y_k}, \ \ \|\mathbf{x}\|_{\mathrm{a}} = \langle \mathbf{x}, \mathbf{x} \rangle_{\mathrm{a}}^{1/2}, \ \ d_{\mathrm{a}}(\mathbf{x}, \mathbf{y}) = \|\mathbf{x} \ominus \mathbf{y}\|_{\mathrm{a}}, \tag{S1}$$

where $\mathbf{x} \ominus \mathbf{y} = \mathbf{x} \oplus \mathbf{y}^{-1}$ is the perturbation difference operation and $\mathbf{y}^{-1} = C[y_1^{-1}, \ldots, y_D^{-1}]$ is the inverse of $\mathbf{y}$.

The clr transformation is defined as $\mathrm{clr}(\mathbf{x}) = \ln\{\mathbf{x}/\mathrm{g}_m(\mathbf{x})\}$, where $\mathrm{g}_m(\mathbf{x}) = (x_1 \ldots x_D)^{1/D}$ is the geometric mean of $\mathbf{x}$. The elements of $\mathrm{clr}(\mathbf{x})$ add to zero. Thus, clr maps $\mathcal{S}^D$ into a subspace of $\mathbb{R}^D$. Next, let $\mathbf{w}_1, \ldots, \mathbf{w}_D$ be compositions obtained by exponentiating $D$ standard unit vectors that form an orthonormal basis for $\mathbb{R}^D$ and applying closure. Hence, $\mathbf{w}_j$ is of the form $C[1, \ldots, e, \ldots, 1]$, where the number $e$ is in column $j$ and 1 is in all other columns. For $\mathbf{x} \in \mathcal{S}^D$, we can write

$$\mathbf{x} = \ln \frac{x_1}{\mathrm{g}_m(\mathbf{x})} \odot \mathbf{w}_1 \oplus \ldots \oplus \ln \frac{x_D}{\mathrm{g}_m(\mathbf{x})} \odot \mathbf{w}_D,$$

implying that the clr transformation provides coefficients for the generating set $\{\mathbf{w}_1, \ldots, \mathbf{w}_D\}$ for $\mathcal{S}^D$. If $\mathbf{u} = \mathrm{clr}(\mathbf{x})$, then $\mathbf{x} = \mathrm{clr}^{-1}(\mathbf{u}) = C[\exp(u_1), \ldots, \exp(u_D)]$ is the inverse transformation. Although the clr transformation uses $D$ coordinates, these coordinates are linearly dependent. As a result, the covariance matrix of clr-transformed data is singular, and probability models directly specified for clr coordinates are degenerate in $\mathbb{R}^D$.

The alr transformation is defined as $\mathrm{alr}(\mathbf{x}) = [\ln(x_1/x_D), \ldots, \ln(x_{D-1}/x_D)]$. It maps $\mathcal{S}^D$ into $\mathbb{R}^{D-1}$. Omitting $\mathbf{w}_D$ from the generating set provides a basis $\{\mathbf{w}_1, \ldots, \mathbf{w}_{D-1}\}$ for $\mathcal{S}^D$. Hence, any $\mathbf{x} \in \mathcal{S}^D$ can be written as

$$\mathbf{x} = \ln \frac{x_1}{x_D} \odot \mathbf{w}_1 \oplus \ldots \oplus \ln \frac{x_{D-1}}{x_D} \odot \mathbf{w}_{D-1}.$$

Thus, the alr transformation provides coefficients for this basis representation. Further, if $\mathbf{v} = \mathrm{alr}(\mathbf{x})$, then $\mathbf{x} = \mathrm{alr}^{-1}(\mathbf{v}) = C[\exp(v_1), \ldots, \exp(v_{D-1}), 1]$ is the inverse transformation.

However, the alr transformation has two important limitations. First, the transformed coordinates depend on the choice of denominator component. Different choices of denominator can lead to different coordinate representations and may affect interpretation. Second, the alr transformation does not preserve distances in the simplex. Therefore, Euclidean distances between alr-transformed vectors do not generally correspond to Aitchison distances between the original compositions.

### Appendix 2: Proofs of propositions.

#### A2.1 Proof of proposition 1

*Proof.* Using the density (13), the cdf of the sampling distribution of $F(x|\hat{\boldsymbol{\mu}}, \hat{\boldsymbol{\Lambda}})$ for any fixed $v \in (0,1)$ can be written as

$$P_{\hat{\boldsymbol{\mu}},\hat{\boldsymbol{\Lambda}}}\{F(x \mid \hat{\boldsymbol{\mu}}, \hat{\boldsymbol{\Lambda}}) \leq v\} = \int \cdots \int I\{F(x \mid \hat{\boldsymbol{\mu}}, \hat{\boldsymbol{\Lambda}}) \leq v\} \times \prod_{i=1}^{n} |\boldsymbol{\Lambda}|^{-1/2} g\{(\mathbf{y}_i - \boldsymbol{\mu})\boldsymbol{\Lambda}^{-1}(\mathbf{y}_i - \boldsymbol{\mu})'\}\, d\mathbf{y}_1 \cdots d\mathbf{y}_n. \tag{S2}$$

where

$$F(x|\hat{\boldsymbol{\mu}}, \hat{\boldsymbol{\Lambda}}) = |\boldsymbol{\Lambda}|^{-1/2} \int I\{(\mathbf{y} - \hat{\boldsymbol{\mu}})\hat{\boldsymbol{\Lambda}}^{-1}(\mathbf{y} - \hat{\boldsymbol{\mu}})' \leq x\} g\{(\mathbf{y} - \boldsymbol{\mu})\boldsymbol{\Lambda}^{-1}(\mathbf{y} - \boldsymbol{\mu})'\} d\mathbf{y}. \tag{S3}$$

Let $\boldsymbol{\Lambda}^{1/2}$ be the symmetric square root of $\boldsymbol{\Lambda}$ and $\boldsymbol{\Lambda}^{-1/2}$ be the inverse of $\boldsymbol{\Lambda}^{1/2}$, both obtained using the spectral decomposition of $\boldsymbol{\Lambda}$. Respectively transform $\mathbf{Y}$ and the data $\mathbf{Y}_i$, $i = 1, \ldots, n$ as

$$\mathbf{U} = (\mathbf{Y} - \boldsymbol{\mu})\boldsymbol{\Lambda}^{-1/2},\ \mathbf{U}_i = (\mathbf{Y}_i - \boldsymbol{\mu})\boldsymbol{\Lambda}^{-1/2}, \tag{S4}$$

giving

$$\mathbf{Y} = \mathbf{U}\boldsymbol{\Lambda}^{1/2} + \boldsymbol{\mu},\ \mathbf{Y}_i = \mathbf{U}_i\boldsymbol{\Lambda}^{1/2} + \boldsymbol{\mu}.$$

Let $(\hat{\boldsymbol{\mu}}_u, \hat{\boldsymbol{\Lambda}}_u)$ be the counterpart of $(\hat{\boldsymbol{\mu}}, \hat{\boldsymbol{\Lambda}})$ for the transformed data. From the affine equivariance property (12), we have

$$\hat{\boldsymbol{\mu}}_u = (\hat{\boldsymbol{\mu}} - \boldsymbol{\mu})\boldsymbol{\Lambda}^{-1/2},\ \hat{\boldsymbol{\Lambda}}_u = \boldsymbol{\Lambda}^{-1/2}\hat{\boldsymbol{\Lambda}}\boldsymbol{\Lambda}^{-1/2},$$

implying that the squared Mahalanobis distance is invariant to the transformation, i.e.,

$$(\mathbf{Y} - \hat{\boldsymbol{\mu}})\hat{\boldsymbol{\Lambda}}^{-1}(\mathbf{Y} - \hat{\boldsymbol{\mu}})' = (\mathbf{U} - \hat{\boldsymbol{\mu}}_u)\hat{\boldsymbol{\Lambda}}_u^{-1}(\mathbf{U} - \hat{\boldsymbol{\mu}}_u)'.$$

Upon applying the transformation (S4) to the integral (S2) and making the above substitution, the integral becomes

$$P_{\hat{\boldsymbol{\mu}},\hat{\boldsymbol{\Lambda}}}\{F(x|\hat{\boldsymbol{\mu}}, \hat{\boldsymbol{\Lambda}}) \leq v\} = \int \ldots \int I\{F(x|\hat{\boldsymbol{\mu}}, \hat{\boldsymbol{\Lambda}}) \leq v\} \prod_{i=1}^{n} g(\mathbf{u}_i\mathbf{u}_i')d\mathbf{u}_1 \ldots d\mathbf{u}_n,$$

where

$$F(x|\hat{\boldsymbol{\mu}}, \hat{\boldsymbol{\Lambda}}) = \int I\{(\mathbf{u} - \hat{\boldsymbol{\mu}}_u)\hat{\boldsymbol{\Lambda}}_u^{-1}(\mathbf{u} - \hat{\boldsymbol{\mu}}_u)' \leq x\} g(\mathbf{u}\mathbf{u}')d\mathbf{u}.$$

This integral does not depend on $(\boldsymbol{\mu}, \boldsymbol{\Lambda})$, establishing the result. □

#### A2.2 Proof of proposition 2

*Proof.* From (13), the likelihood function for $(\boldsymbol{\mu}, \boldsymbol{\Lambda})$ based on data $\mathbf{y}_1, \ldots, \mathbf{y}_n$ is

$$L(\boldsymbol{\mu}, \boldsymbol{\Lambda}) = \prod_{i=1}^{n} |\boldsymbol{\Lambda}|^{-1/2} g\{(\mathbf{y}_i - \boldsymbol{\mu})\boldsymbol{\Lambda}^{-1}(\mathbf{y}_i - \boldsymbol{\mu})'\}. \tag{S5}$$

Suppose the data are transformed as $\mathbf{u}_i = \mathbf{y}_i\mathbf{A} + \mathbf{b}$, $i = 1, \ldots, n$, as stipulated in (12) for affine equivariance. The likelihood function based on the transformed data can be written as

$$L(\boldsymbol{\mu}_u, \boldsymbol{\Lambda}_u) = \prod_{i=1}^{n} |\boldsymbol{\Lambda}_u|^{-1/2} g\{(\mathbf{u}_i - \boldsymbol{\mu}_u)\boldsymbol{\Lambda}_u^{-1}(\mathbf{u}_i - \boldsymbol{\mu}_u)'\}, \tag{S6}$$

where $\boldsymbol{\mu}_u = \boldsymbol{\mu}\mathbf{A} + \mathbf{b}$ and $\boldsymbol{\Lambda}_u = \mathbf{A}'\boldsymbol{\Lambda}\mathbf{A}$. Maximization of (S5) gives ML estimator $(\hat{\boldsymbol{\mu}}, \hat{\boldsymbol{\Lambda}})$ of $(\boldsymbol{\mu}, \boldsymbol{\Lambda})$, whereas maximization of (S6) gives ML estimator $(\hat{\boldsymbol{\mu}}_u, \hat{\boldsymbol{\Lambda}}_u)$ of $(\boldsymbol{\mu}_u, \boldsymbol{\Lambda}_u)$. However, from the invariance property of ML estimators, we have $(\hat{\boldsymbol{\mu}}_u, \hat{\boldsymbol{\Lambda}}_u) = (\hat{\boldsymbol{\mu}}\mathbf{A} + \mathbf{b}, \mathbf{A}'\hat{\boldsymbol{\Lambda}}\mathbf{A})$. From (12), this establishes the desired affine equivariance property of the ML estimators. □

##### A2.3 Proof of proposition 4

*Proof.* The tolerance region that goes through point $\mathbf{z}_0$ is an ellipsoid of the form

$$T = \{\mathbf{z}_0 \in \mathbb{R}^d : \mathrm{MD}_{\mathbf{z}_0}(\hat{\boldsymbol{\mu}}, \hat{\boldsymbol{\Lambda}}) \leq c_0(d, p_0, \alpha, n)\}, \tag{S7}$$

where the tolerance factor $c_0$ is chosen to satisfy the probability requirement

$$1 - \alpha = P_{\hat{\boldsymbol{\mu}}, \hat{\boldsymbol{\Lambda}}} \left\{ P_{\mathbf{Z}}\big(\mathbf{Z} \in T \mid \hat{\boldsymbol{\mu}}, \hat{\boldsymbol{\Lambda}}\big) \geq p_0 \right\} \tag{S8}$$

We can write (S8) as

$$1 - \alpha = P_{\hat{\boldsymbol{\mu}}, \hat{\boldsymbol{\Lambda}}} \left\{ F(c_0 | \hat{\boldsymbol{\mu}}, \hat{\boldsymbol{\Lambda}}) \geq p_0 \right\} \tag{S9}$$

Let $F(c_0|\hat{\boldsymbol{\mu}}, \hat{\boldsymbol{\Lambda}}) = U$. Then we can write (S9) as

$$1 - \alpha = F\,(U \geq p_0)\; = 1 - P(U \leq p_0) \tag{S10}$$

and (S10) as

$$\alpha = P(U \leq p_0) = F_U(p_0) \tag{S11}$$

and (S11) as

$$F_U^{-1}(\alpha) = p_0$$

□

#### Appendix 3: Some basic compositional data analysis concepts

##### A3.1 Measures of center and dispersion

For data $x_{ij}$, $j = 1, \ldots, D$, $i = 1, \ldots, n$ on $D$-part compositions from $n$ subjects, the measures of central tendency and dispersion are respectively given by the *geometric center* and the *variation matrix* and *total variance.*[1] The geometric center $\mathrm{cen}(\mathbf{X}) = C[\hat{g}_1, \ldots, \hat{g}_D]$ is the closed column-wise sample geometric mean of the data, where $\hat{g}_j = \prod_{i=1}^{n}(x_{ij})^{1/n}$. The variation matrix $\mathbf{T} = (t_{jk})$ is a $D \times D$ matrix whose general element

$$t_{jk} = \mathrm{var}\left(\frac{1}{\sqrt{2}} \ln \frac{x_j}{x_k}\right) = \frac{1}{n} \sum_{i=1}^{n} \left(\ln \frac{x_{ij}}{x_{ik}} - \ln \frac{\hat{g}_j}{\hat{g}_k}\right)^2$$

is the sample variance of logratio of parts $j$ and $k$ calculated using the geometric center. A small value for $t_{jk}$ indicates that the ratio $x_j/x_k$ is nearly constant, suggesting a strong relationship between the parts $j$ and $k$. The matrix $\mathbf{T}$ has zeros on the diagonal. Further, since all parts of the composition share a common scale, the logratio variances can be added up to get a measure of overall variability in the data: $\text{totvar}(\mathbf{X}) = \frac{1}{2D}\sum_{j,k=1}^{D} t_{jk}$.

### Appendix 4: Additional results for reference regions for microbiome data

#### A4.1 Three-part subcomposition

##### A4.1.1 Location and covariance estimates:

The main article provides estimates for location ($\boldsymbol{\mu}$) and covariance ($\boldsymbol{\Sigma}$) of ilr coordinates using the classical and SD method. Here we provide the estimates using the other methods.
MCD method:

$$\mathbf{M}_z = \begin{bmatrix}1.077 & 0.665\end{bmatrix}, \ \mathbf{V}_z = \begin{bmatrix} 0.272 & -0.035 \\ -0.035 & 0.635 \end{bmatrix}.$$

S-estimation method:

$$\mathbf{M}_z = \begin{bmatrix}1.033 & 0.714\end{bmatrix}, \ \mathbf{V}_z = \begin{bmatrix} 0.253 & -0.082 \\ -0.082 & 0.571 \end{bmatrix}.$$

##### A4.1.2 Normality assessment for three part composition

Figure S1 a display the normal Q–Q plots for each ilr coordinate and chi-square Q–Q plots for the multivariate normal distribution. Since the points in the plots more or less form a straight line, the assumption of normality appears quite reasonable for these data.

#### A4.2 Eight-part composition

##### A4.2.1 Geometric center and dispersion matrix

$$\text{cen}(\mathbf{X}) = \begin{bmatrix}0.1761 & 0.4356 & 0.0003 & 0.1931 & 0.0026 & 0.0765 & 0.0010 & 0.1147\end{bmatrix}$$

$$\mathbf{T} = \begin{bmatrix}
0.0000 & 0.0946 & 1.3607 & 0.8284 & 1.8961 & 0.7935 & 1.4148 & 0.3650 \\
0.0946 & 0.0000 & 1.4238 & 0.8674 & 1.9277 & 0.5904 & 1.5168 & 0.2547 \\
1.3607 & 1.4238 & 0.0000 & 0.5326 & 0.9059 & 2.0987 & 0.4371 & 2.0961 \\
0.8284 & 0.8674 & 0.5326 & 0.0000 & 0.9677 & 1.3933 & 0.5094 & 1.5629 \\
1.8961 & 1.9277 & 0.9059 & 0.9677 & 0.0000 & 2.7725 & 0.6225 & 2.5926 \\
0.7935 & 0.5904 & 2.0987 & 1.3933 & 2.7725 & 0.0000 & 2.2506 & 1.0011 \\
1.4148 & 1.5168 & 0.4371 & 0.5094 & 0.6225 & 2.2506 & 0.0000 & 2.1937 \\
0.3650 & 0.2547 & 2.0961 & 1.5629 & 2.5926 & 1.0011 & 2.1937 & 0.0000
\end{bmatrix}$$

##### A4.2.2 Location and covariance estimates:

Normal model:

$$\bar{\mathbf{Z}} = \begin{bmatrix}2.1351 & 3.4438 & -3.8902 & 2.4551 & -1.8082 & 1.5882 & -3.3240\end{bmatrix}$$

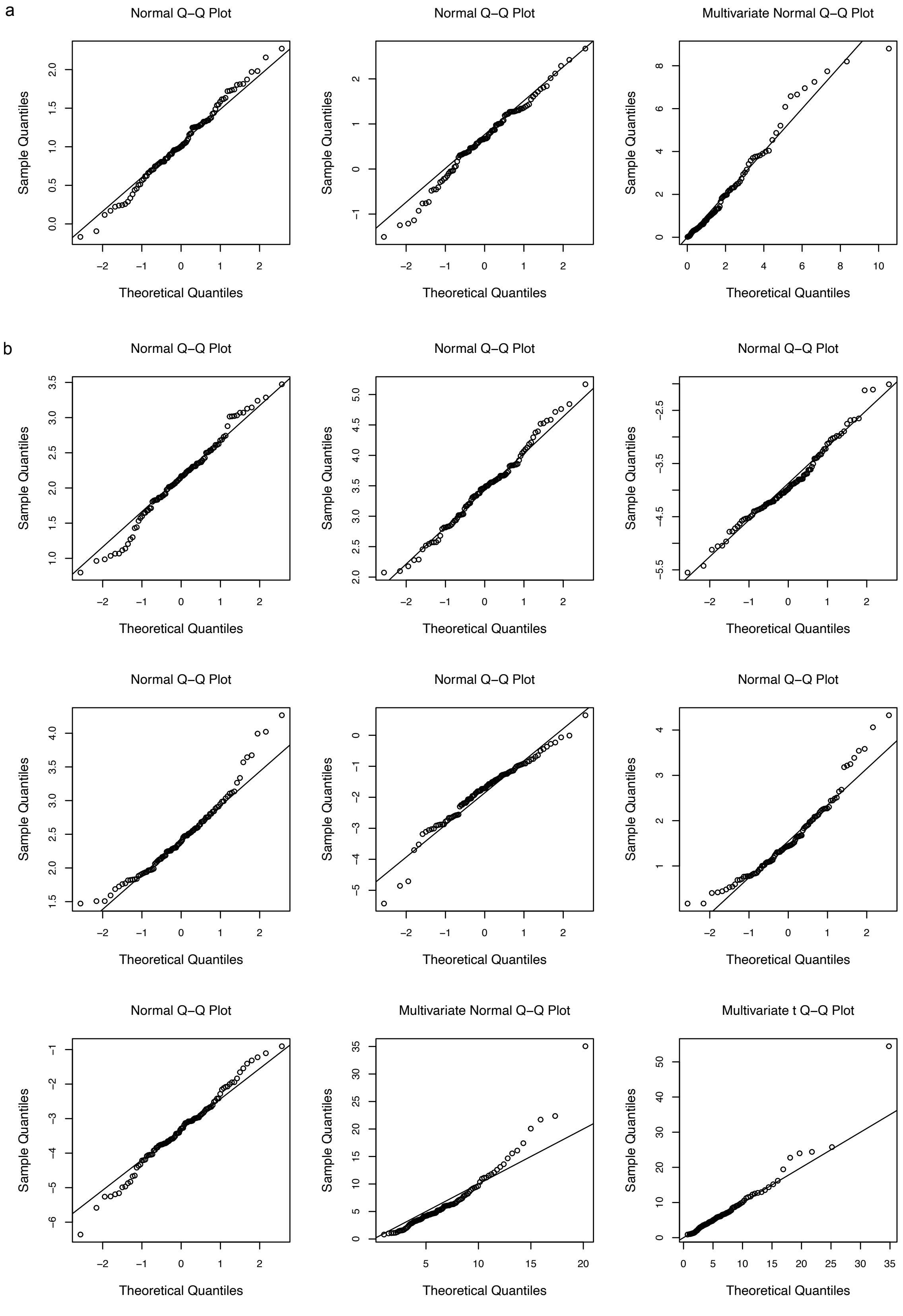


Figure S1: Normal Q–Q plots for each ilr coordinate and chi-square Q–Q plots for the multivariate distributions. **a** Results for the 3-part composition, where the chi-square Q–Q plot is based on the multivariate normal assumption. **b** Results for the 8-part composition, where the chi-square Q–Q plots are shown for both the multivariate normal and multivariate $t$ distributions.

$$\mathbf{S}_z = \begin{bmatrix} 0.3349 & 0.3265 & -0.1955 & -0.0852 & -0.3814 & 0.1181 & -0.4498 \\ 0.3265 & 0.4178 & -0.2707 & -0.1389 & -0.4882 & 0.2242 & -0.6004 \\ -0.1955 & -0.2707 & 0.4829 & 0.1525 & 0.2660 & -0.2390 & 0.4209 \\ -0.0852 & -0.1389 & 0.1525 & 0.3150 & 0.1076 & -0.0759 & 0.2762 \\ -0.3814 & -0.4882 & 0.2660 & 0.1076 & 1.0431 & -0.4788 & 0.7307 \\ 0.1181 & 0.2242 & -0.2390 & -0.0759 & -0.4788 & 0.7183 & -0.3607 \\ -0.4498 & -0.6004 & 0.4209 & 0.2762 & 0.7307 & -0.3607 & 1.0968 \end{bmatrix}$$

$t$ model:

$$\boldsymbol{\mu} = \begin{bmatrix} 2.1224 & 3.4368 & -3.9079 & 2.3971 & -1.7945 & 1.5338 & -3.3278 \end{bmatrix}$$

$$\boldsymbol{\Lambda} = \begin{bmatrix} 0.3498 & 0.3390 & -0.1824 & -0.1108 & -0.4095 & 0.1014 & -0.4783 \\ 0.3390 & 0.4203 & -0.2510 & -0.1498 & -0.4901 & 0.2024 & -0.6014 \\ -0.1824 & -0.2510 & 0.4710 & 0.1751 & 0.2299 & -0.2047 & 0.3872 \\ -0.1108 & -0.1498 & 0.1751 & 0.2863 & 0.1041 & -0.1109 & 0.2819 \\ -0.4095 & -0.4901 & 0.2299 & 0.1041 & 1.0551 & -0.4746 & 0.6990 \\ 0.1014 & 0.2024 & -0.2047 & -0.1109 & -0.4746 & 0.6496 & -0.3509 \\ -0.4783 & -0.6014 & 0.3872 & 0.2819 & 0.6990 & -0.3509 & 1.0792 \end{bmatrix}$$

##### A4.2.3 Normality assessment for eight part composition:

Figure S1 display the normal Q–Q plots for each ilr coordinate and chi-square Q–Q plots for the multivariate normal and $t$ distributions. In marginal Q–Q plots the points more or less form a straight line, indicating a reasonable fit. However, when looking at the multivariate normal Q-Q plot, we observe heavier tails, suggesting potential deviations from normality in the joint distribution. On the other hand, the multivariate t Q-Q plot shows that the assumptions are generally reasonable, with the data points aligning well with the $t$ distribution except for one outlier point.

#### A4.3 Diagnostic plots for liver cirrhosis and type 2 diabetes data

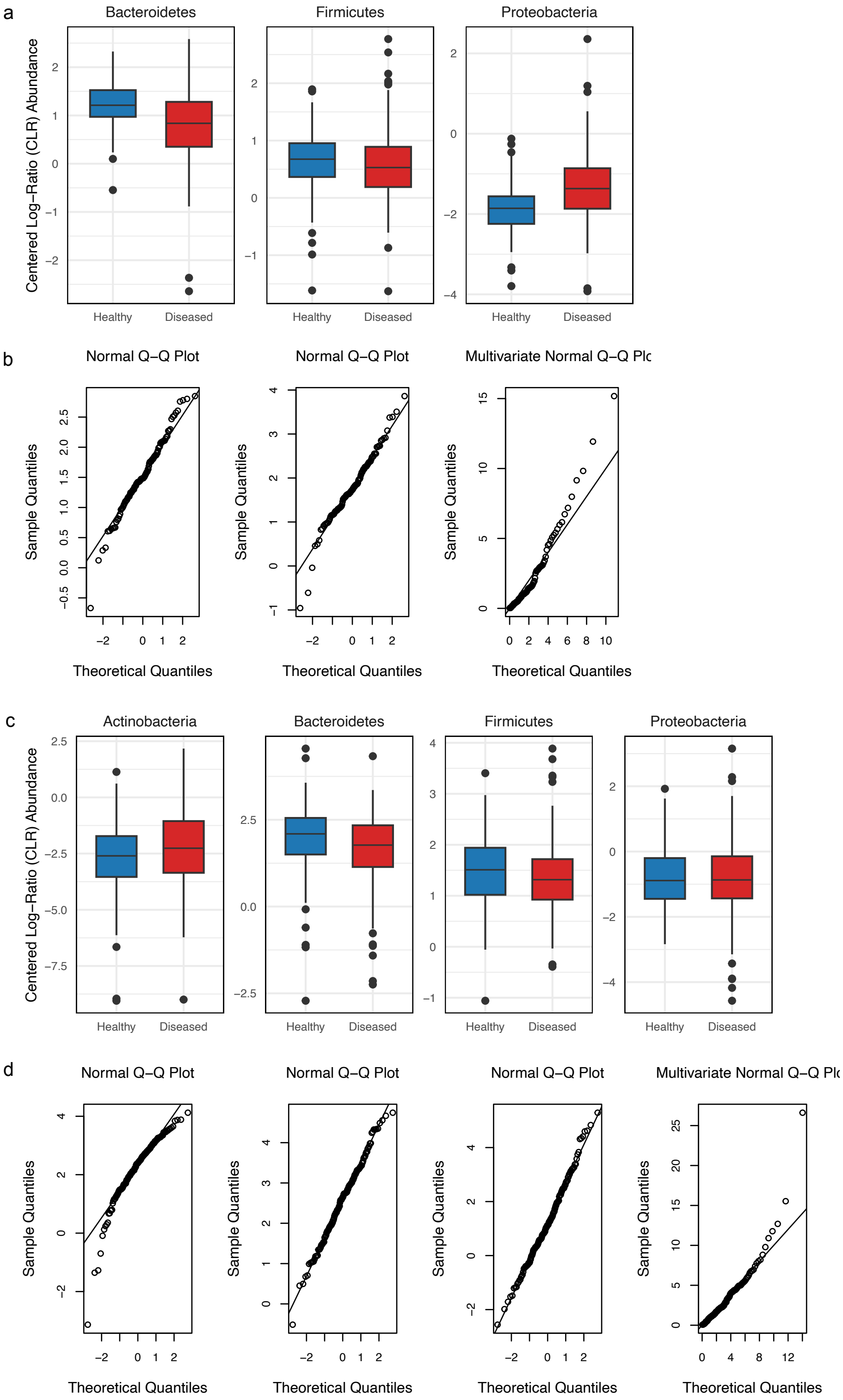

Figure S2: **a,c** Centered log-ratio (CLR)-transformed boxplots used to visually assess distributional differences between healthy and diseased groups for the liver cirrhosis and type 2 diabetes studies, respectively. **b,d** Normal Q–Q plots for each ilr coordinate and chi-square Q–Q plots for the corresponding multivariate distributions in the liver cirrhosis and type 2 diabetes studies, respectively.

#### References


[1] Vera Pawlowsky-Glahn, Juan José Egozcue, and Raimon Tolosana-Delgado. *Modeling and Analysis of Compositional Data.* John Wiley & Sons, 2015.
